# Spatially-embedded recurrent neural networks reveal widespread links between structural and functional neuroscience findings

**DOI:** 10.1101/2022.11.17.516914

**Authors:** Jascha Achterberg, Danyal Akarca, DJ Strouse, John Duncan, Duncan E Astle

**Author notes:** Co-lead authors (Jascha Achterberg, Danyal Akarca). Co-senior authors (John Duncan, Duncan E Astle). **Corresponding authors:** Jascha Achterberg, Danyal Akarca.

## Abstract

Brain networks exist within the confines of resource limitations. As a result, a brain network must overcome metabolic costs of growing and sustaining the network within its physical space, while simultaneously implementing its required information processing. To observe the effect of these processes, we introduce the spatially-embedded recurrent neural network (seRNN). seRNNs learn basic task-related inferences while existing within a 3D Euclidean space, where the communication of constituent neurons is constrained by a sparse connectome. We find that seRNNs, similar to primate cerebral cortices, naturally converge on solving inferences using modular small-world networks, in which functionally similar units spatially configure themselves to utilize an energetically-efficient mixed-selective code. As all these features emerge in unison, seRNNs reveal how many common structural and functional brain motifs are strongly intertwined and can be attributed to basic biological optimization processes. seRNNs can serve as model systems to bridge between structural and functional research communities to move neuroscientific understanding forward.

## INTRODUCTION

As they develop, brain networks learn to achieve objectives, from simple functions like autonomic regulation, to higher-order processes like solving problems. Many stereotypical features of networks are downstream consequences of resolving challenges and trade-offs they face, across their lifetime (Fair et al., 2009; Kaiser, 2017) and evolution (Bosman & Aboitiz, 2015; Heuvel et al., 2016; Hiratani & Latham, 2022). One example is the optimization of functionality within resource constraints; all brain networks must overcome metabolic costs to grow and sustain the network in physical space, whilst simultaneously optimizing that network for information processing. This trade-off shapes all brains within and across species, meaning it could be why many brains converge on similar organizational solutions (Heuvel et al., 2016). As such, the most basic features of both brain organization and network function – like its sparse and small-world structure, functional modularity, and characteristic neuronal tuning curves – might arise in unison as a result of this basic optimization problem.

Our understanding of how the brain’s structure and function interact largely comes from observing differences in brain structure, such as across individuals (Mišić et al., 2016) or following brain injury (Smith et al., 2022), and then systematically linking these differences to brain function or behavioral outcomes. But how do these relationships between structure, function and behavior emerge *in the first place*? To address this question, we need to be able to manipulate experimentally how neural networks form, *as they learn to achieve behavioral objectives*, in order to establish the causality of these relationships. Computational models allow us to do this. They have shown that network modularity can arise through the spatial cost of growing a network (Kaiser & Hilgetag, 2004), how orthogonal population dynamics can arise purely through optimizing task performance (Mante et al., 2013) and how predictive coding can arise through limiting a brain’s energy usage (Ali et al., 2021). But we have yet to incorporate both the brain’s anatomy and function into a single coherent model, allowing a network to dynamically trade-off its different structural, functional, and behavioral objectives in real time.

To achieve this, we introduce spatially-embedded recurrent neural networks (seRNNs). An seRNN is optimized to solve a task, making decisions to achieve functional goals. However, as it learns to achieve these goals its constituent neurons face the kind of resource constraints experienced within biological networks. Neurons must balance their finite resources to grow or prune connections, whilst the overall network attempts to optimize intra-network communication and behavioral performance. By allowing seRNNs to dynamically manage both their structural and functional objectives simultaneously, while they learn to behave, multiple simple and complex hallmarks of biological brains naturally emerge.

## RESULTS

### How to spatially embed a recurrent neural network

We created an optimization process for a general recurrent neural network (RNN). This optimization allows flexible trade-offs between improving task performance within the resource constraints of existing in a biophysical space. Specifically, this biophysical space acts as a prior, incentivizing the network to prune long distance connections in 3D physical space while supporting its intra-network communication of signals.

In a canonical supervised RNN, all the network’s trainable parameters are optimized to minimize the difference between the predicted value and correct value. To achieve this, we define a task loss function (*L*) which defines the prediction error to be minimized to optimize task performance. To produce a network that generalizes well to unseen data, we can add a regularization term.

Regularization incentivizes networks to converge on sparse solutions and is commonly applied to neural networks in general (Hardt & Recht, 2022) and neuroscientific network models (Kietzmann et al., 2019; Yang et al., 2019). For a regularized network, the loss function becomes a combination of both the task loss and the regularization loss. One example of a commonly applied regularization is the L1 regularization, which is also used in LASSO regression (Tibshirani, 1996) and incentivizes the network to maximize task performance while concurrently minimizing the sum of all absolute weights in the neural network. If we want to regularize the recurrent weight matrix (*W*) with the dimensions *m* × *m* where *m* is number of units in the recurrent layer, the loss function would be:

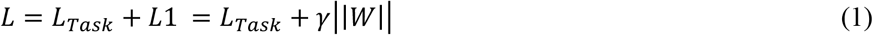

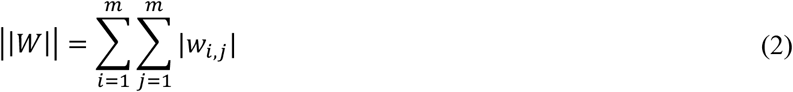

An RNN with this loss function would learn to solve the task with a sparse weight matrix, where *γ* would determine the extent to which the network is forced to converge on a sparse solution. This parameter is called the regularization strength.

Unlike regular RNNs, real brain networks are embedded in a physical space (Barthélemy, 2011; Bassett & Stiso, 2018; Bullmore & Sporns, 2012). To simulate the pressures caused by existing in a biophysical space, we manipulated the regularization term. We hypothesized that incorporating constrains that appear common to any biological neural system, we could test whether these local constraints are sufficient to drive a network architecture that more closely resembles observed brain networks. Specifically, we included spatial constraints in two forms – Euclidean and network communication – that we argue are integral to any realistic neural network.

The Euclidean embedding makes sure that neurons are not mathematically abstract computational nodes, but instead exist within a physical locality in which space is meaningful. This is important, because the physical distance between neurons is a critical influence on their connectivity (Akarca et al., 2021; Bullmore & Sporns, 2012; Song et al., 2014). To implement this, we embed units within a 3D space, such that each unit has a corresponding x, y, and z coordinate. Using these coordinates, we can generate a Euclidean distance matrix which describes the physical distance between each pair of nodes (**Figure 1a**). This allows to minimize weights multiplied by their Euclidean distance (*d*_*i, j*_), thereby incentivizing the network to minimize (costly) long distance connections. The element-wise matrix multiplication is denoted with the Hadamard product ⊙. Adding this to our optimization term gives us:

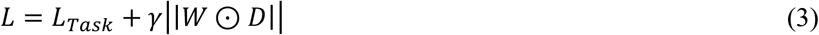

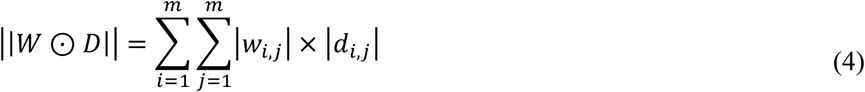

The above formalization provides a spatial context for RNN training. However, a further critical feature found in physical neural systems is that their communication dynamics are constrained (Avena-Koenigsberger et al., 2018; Laughlin & Sejnowski, 2003). That is, neurons communicate with each other as a function of how their signals propagate through the network to their neighbors (and their neighbors’ neighbors, and so on). Intra-network communication becomes a central factor in the pruning process of networks, as weaking connections in a fully connected network will always reduce how well information can flow through a network (Crofts & Higham, 2009; Griffa et al., 2022; Seguin, Mansour L, et al., 2022). So, network communicability is an important prior to guide a network’s pruning process while it learns to solve a task.

**Figure 1.**
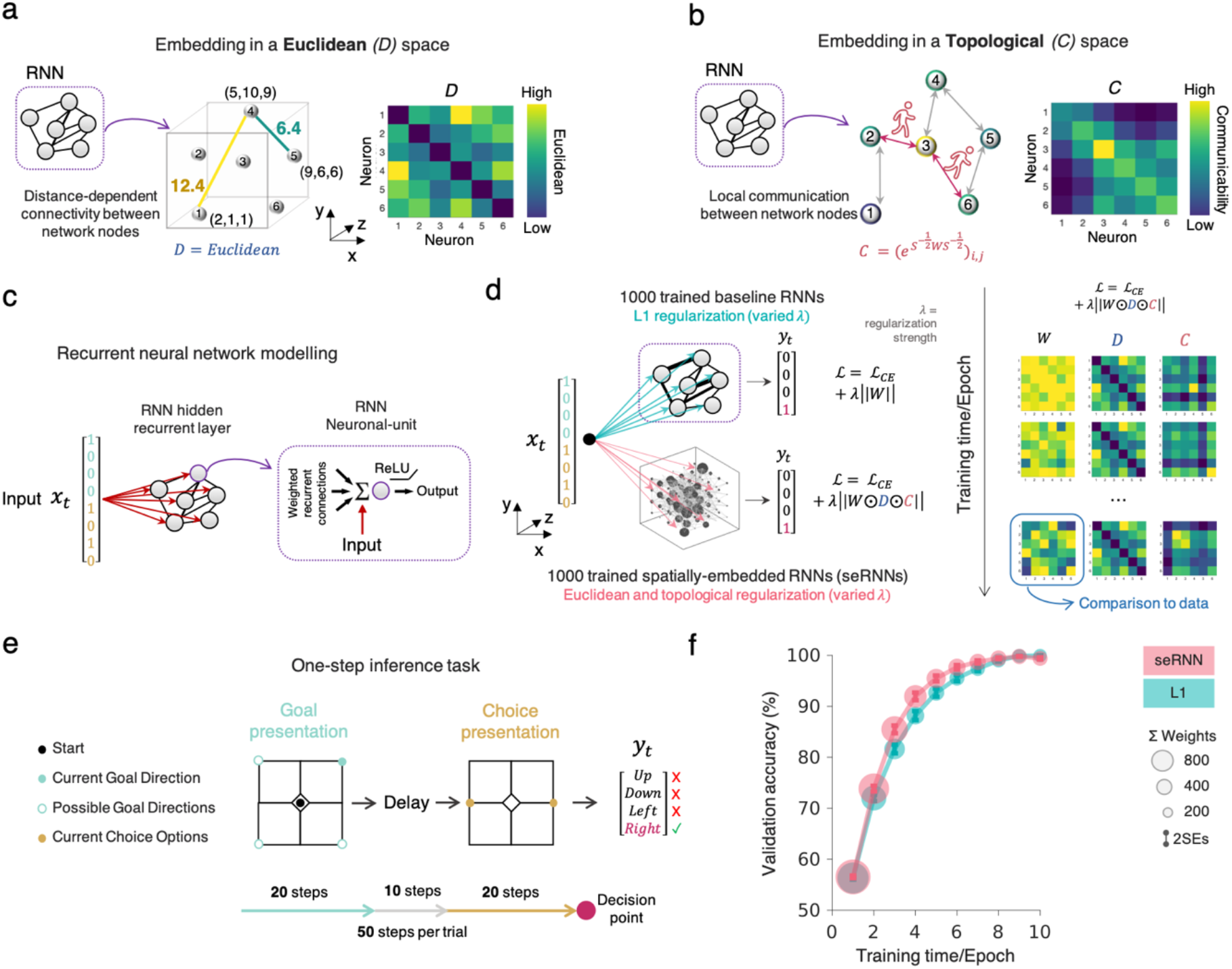
Task structure and spatially-embedded recurrent neural networks (seRNNs). **a** We embed RNN in Euclidean space by assigning each of the 100 units a location on an evenly spaced 5x5x4 grid. We show a schematic demonstration of a small six-node network and show an analogous Euclidean distance matrix. **b** We also provided topological communication constraints to the RNN to guide the pruning process towards good intra-network communication. This topological constrain is the weighted communicability measure, which is equivalent to an infinite random walk across the network on a weighted connectivity matrix. The weighted communicability term of the matrix is shown for the same simple six-node network. **c** A recurrent neural network (RNN) was trained to complete a task. Each unit of the 100-unit RNN was provided with a vector of input information at the relevant step points. Each neural unit consisted of a ReLU activation function, which also simultaneously took all weighted recurrent connections and provided a non-linear output. **d** For the main part of the study, we trained 1000 standard L1-regularized RNNs as a baseline comparison to 1000 spatially-embedded RNNs (seRNNs) which entails both the Euclidean and topological constraints. The training was undertaken such that Euclidean distances and weighted communication were minimized. To the right we provide a demonstration on a simple network how the *W, D* and *C* matrix all change over training epochs. The distance matrix is static due to fixed locations of units, but the communicability matrix changes as the structure of the network (represented by the weight matrix) changes. **e** The one-step inference task solved by networks consists of an initial period of 20 steps in which a goal is presented in one of four locations on the grid space: top left, top right, bottom left, or bottom right. This is depicted in light blue in the top right corner of the grid. Subsequently, there is a 10-step delay period in which this goal location must be kept in memory. Then two choice options are provided for 20 steps. Using the goal location information from before, agents must choose the choice option which is closer to the goal. In this example, as a left and right choice option are provided, the correct decision at the end of this 20-step choice presentation period is to select right. **f** The validation accuracy of all converging neural networks is shown across L1 RNNs (blue) and seRNNs (pink), showing the equivalent performance is achieved on the one-step inference task.

How can we model this communication constraint? In the above equation, despite accounting for the spatial location, topological communication has no influence on connection updates. We can impose the influence of communication via a *network communicability* term (C; Crofts & Higham, 2009), which computes the extent to which, under a particular network topology, any two nodes are likely to communicate both directly and indirectly over time (**Figure 1b**). Now taking this topological communication into account, we get the following loss function:

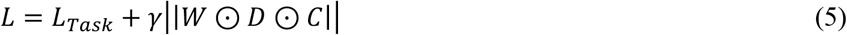

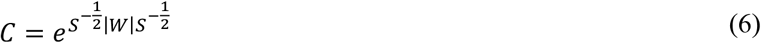

**Supplementary Figures 1-6** provides a walk-through explanation of how this term works and expand on the logic of how constraining the network’s topology can serve as a prior for intra-network communication in sparse networks. **Supplementary Figure 6** specifically highlights the role that communicability has within the network optimization process in terms of both global (i.e., the whole network) and local (i.e., connections within a network) communication. Note in **Equation 6** that *S* is a diagonal matrix with the degree of unit *i* (*deg*_*i*_) on the diagonal (i.e., the node strength) which simply acts as a normalization term preventing any one single edge having undue influence (Crofts & Higham, 2009). This is explained in **Supplementary Figure 4, 5**.

We now have constructed a loss function bringing together task control with both spatial and communication network constraints into a single optimization term. To understand how this spatial embedding impacts a network’s optimization process, we set up a population of 2000 networks. All networks are standard Recurrent Neural Networks, with 100 units in the hidden layer and Rectified Linear Unit (ReLU) activation functions (**Figure 1c**). They are optimized using Adam for 10 epochs. Half of the networks (1000 networks) are regularized using the custom seRNN regularizer described above, which we term seRNNs. For this, we spatially embed the 100 units of the recurrent layer by assigning them locations on a 5x5x4 grid, which defines the Euclidean *D* term. The other half (1000 networks) are regularized using a standard L1 regularizer as we outlined above (the baseline model). In both cases, the regularizer is only applied to the hidden recurrent layer of the network and the regularization strength is systematically varied within each subgroup of networks to cover a wide spectrum of regularization strength which is matched across subgroups (**Figure 1d**, see **Methods; *Regularization strength setup and network selection***). All our networks start strongly connected (using an orthogonal initialization) and learn through a guided pruning process (Ducharme et al., 2016; Huttenlocher, 1979; Tamnes et al., 2017). We trained all RNNs on a simple one-choice inference task which required networks to develop two fundamental functions of recurrent networks: Remembering and integration of information (**Figure 1e**, see **Methods; *Task paradigm***). On an abstract level, networks needed to first store a stimulus, integrate it with a second stimulus, and make a predefined correct choice. Our task setup can also be interpreted as a simple one-choice maze inference task. In this interpretation, networks were trained to observe a goal location in one of four possible locations (the corners of a 3x3 grid), before a short delay period in which the goal location is removed and needs to be remembered. After the delay, two possible directions are given as choice options. The choice option closer to the goal location is the correct target. Both the task input and choice options are One-Hot encoded, with a low level of Gaussian noise added to the task inputs. When training all networks, we find that both types of networks successfully manage to learn the task with high accuracy (both can achieve > 95% average accuracy; **Figure 1f**).

We tested whether and how spatially embedding the RNNs influences the structure and function of the networks which successfully solve the task (> 90% task accuracy, see **Methods; *Regularization strength setup and network selection***). Specifically, using L1 networks as a baseline, we tested whether seRNNs show features we commonly observe in the brain – especially in primate cerebral cortices. Initially we look at purely structural motifs, including modularity (Bertolero et al., 2015; Park & Friston, 2013; Sporns & Betzel, 2016) and small-worldness (Bassett & Bullmore, 2017; Sporns & Zwi, 2004), and then progress to testing whether functionally similar units cluster in space, as they usually do in brains (Bassett & Bullmore, 2017; Sporns & Zwi, 2004). Following this, we tested whether the impact of the spatial embedding goes beyond the structural and functional organization of networks and influences how a network’s units work together to achieve high task performance. Task solving networks in the brain often implement an energy efficient *mixed selective code* (Johnston et al., 2020; Rigotti et al., 2013). We wanted to see whether this would occur seRNNs. In short, we wanted to test if established organization properties of complex brain networks arise when we impose local biophysical constraints.

### Modular small-world recurrent networks emerge from Euclidean and communication constraints

We first investigated the structural topology of our trained networks. In **Figure 2a** we show a representative example of an seRNN network depicted in the 3D Euclidean space it was trained in, showing sparse connectivity. In accordance with the training regime, both L1 and seRNN networks develop a sparse connectome due to the objective function minimizing weights. In both cases this sparsity increases over the course of the training time (**Figure 2b, left**). However, seRNN networks - due to being embedded in a physical space - penalize their weights such that connections which are further away tend to become weaker. This is commonly found in empirical brain networks across species and scales (e.g., Betzel et al., 2018) (**Figure 2b, right**).

**Figure 2.**
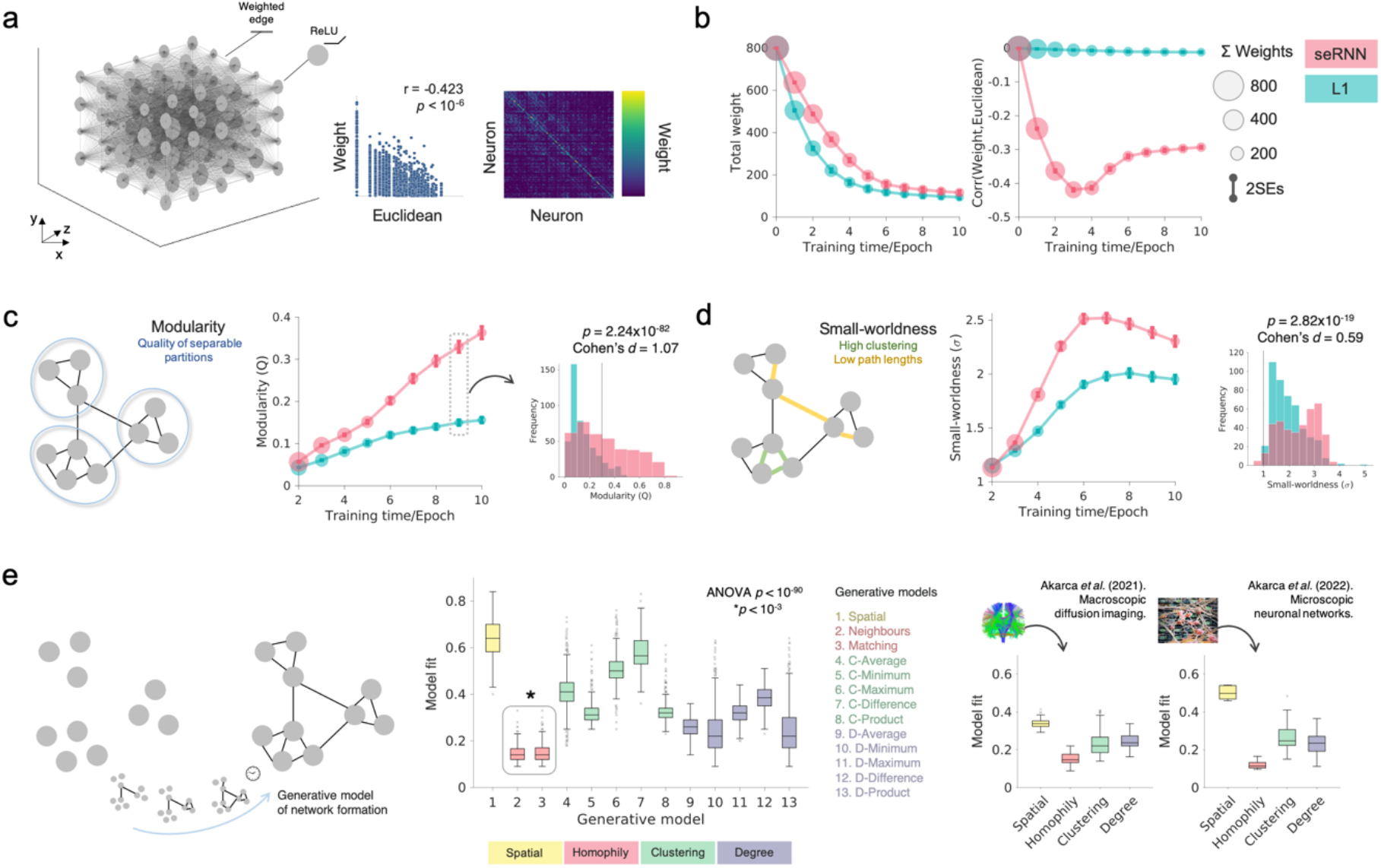
Spatially embedded recurrent neural networks (seRNNs) exhibit a brain-like structural topology. **a** Left, an example of a representative seRNN network in the 3D space in which it was trained. This network was taken from epoch 9 at a regularization of 0.08 and is the network used for visualizations for the rest of this paper. Middle, we show the negative relationship between the connection weights of seRNN versus the Euclidean distances of the connections. Right, we show the weight matrix of this seRNN. **b** Left, weights decline for both L1 and seRNN networks over the course of epochs/training. Right, seRNNs have a negative correlation between weight and Euclidean distance over the course of epochs/training, but in L1 networks there is no relationship between weights and Euclidean distance. These results are across all networks. **c** Left, a schematic illustration of the concept of modularity in networks. While both L1 and seRNN networks exhibit increasing modularity over epochs/training, there is a consistently greater modularity in seRNNs compared to L1 networks. Right, we show very large (Cohen’s *d* = 1.07) statistical differences in modularity distributions for functioning (validation accuracy ≥ 90%) epoch 9 networks in L1 and seRNN networks. **d** Left, a schematic illustration of the concept of small-worldness in networks. While both L1 and seRNN networks exhibit a similar trajectory shape of small-worldness over epochs/training, there is a consistently greater small-worldness in seRNNs compared to L1 networks. Right, we show moderate-to-large (Cohen’s *d* = 0.59) statistical differences in small-worldness distributions for functioning epoch 9 networks in L1 and seRNN networks. **e** For a range of generative network models, we present the model fit of the top performing simulations fit (see **Methods; *Voronoi tessellation parameter fitting procedure***) to seRNNs. Results show that homophily models achieve the best model fits. These findings are congruent with published data from adolescent whole-brain diffusion-MRI connectomes (bottom, left) and high-density functional neuronal networks at single-cell resolution (bottom, right).

We next investigated two key topological characteristics that are commonly found in empirical brain networks across spatial scales and proposed to facilitate brain function: modularity (Bertolero et al., 2015; Park & Friston, 2013; Sporns & Betzel, 2016) and small-worldness (Bassett & Bullmore, 2017; Sporns & Zwi, 2004). Modularity denotes dense intrinsic connectivity within a module but sparse weak extrinsic connections between modules and is commonly measured via a Q statistic (denoting how well sub-communities within the network can be partitioned (Newman, 2006)). In contrast, small-worldness indicates a short average path length between all node pairs, with high local clustering – whereby networks reside somewhere between random and ordered networks (see **Methods; *Topological analysis***). We tested if these properties emerged in both our L1 and seRNN model systems.

Computing modularity Q statistics and small-worldness relative to appropriate null models (see **Methods; *Topological analysis***) shows that seRNNs consistently exhibit both increased modularity (**Figure 2c**) and small-worldness (**Figure 2d**) relative to L1 networks over the course of training.

Differences are smaller initially, but later in training, the effect size for differences in modularity are large (at epoch 9, modularity *p* = 2.24x10^−82^, Cohen’s *d* = 1.07; **Figure 2c, right**) and for small-worldness moderate-to-large (*p* = 2.82 × 10^−19^, Cohen’s *d* = 0.59; **Figure 2d, right**). It is of note that, while there is variability across empirical brain networks depending on species, modality and scale, seRNNs achieve modularity Q statistics within ranges commonly found in empirical human cortical networks (Hilger et al., 2017). Both L1 and seRNNs achieve the technical definition of small-worldness of >1 (Watts & Strogatz, 1998), but seRNNs exhibit a higher value more consistent with empirical networks (for review, see Bassett & Sporns, 2017). **Supplementary Figure 7** shows how the subparts of the regularization interact with the task optimization to shape these structural effects. It is important to note that within the population of seRNNs we find varying degrees of modularity and small-worldness (**Figure 2c, right; Figure 2d, right**) – some seRNNs in fact show lower modularity values than some L1 networks. We will return to this variability in a later section and provide an explanation for how it can be understood.

Beyond observing topology alone, in recent years generative network models (Kaiser & Hilgetag, 2004) have been employed to uncover what wiring rules can be used to best simulate empirical network topology *in silico* (Akarca et al., 2021; Betzel et al., 2016; Vértes et al., 2012). These models work by simulating the formation of networks over time within a 3D space, by probabilistically adding connections. This is determined by a probabilistic wiring equation, which balances the costs of forming connections with a particular topological wiring rule (see **Methods**; ***Generative network models***). Recently, one particularly successful type of wiring rule has been *homophily* – where nodes preferentially wire with other nodes that are similar to themselves in terms of their shared connectivity. These rules have been shown to very effectively simulate the statistical topology of both empirical structural and functional connectivity, across scales and species (Akarca et al., 2021, 2022; Betzel et al., 2016; Carozza et al., 2022; Oldham et al., 2022; Vértes et al., 2012). While it has been observed that homophily resonates with Hebbian-like mechanisms (Akarca et al., 2022; Goulas et al., 2019; Vértes et al., 2012) it still remains unclear *how* or *why* such rules would be implemented in neurobiological networks. To address this, we next tested the possibility that this homophily phenomenon can be explained simply by the spatial and communication constraints intrinsic to biological neural networks under task-control. That is, the fact that homophily mechanisms produce realistic topology is an epiphenomenon of the underlying basic physical constraints that brain networks are under. If so, we should expect that, just as in empirical brain networks, the topology of seRNNs should also be best recapitulated by a homophily generative rule.

By fitting generative network models (see **Methods; *Model fitting***) to functioning seRNNs (**≥**90% validation accuracy) we indeed find that the topology of seRNNs are uniquely best simulated by homophily generative models (One-way ANOVA F(12,2730) = 814.3, *p* < 10^−90^; all Tukey HSD comparisons to other rules, *p* < 3x10^−7^) (**Figure 2e**). This is not the case for L1 networks (one-way ANOVA F(12,3783) = 546.4, *p* < 10^−90^; both Tukey HSD comparisons with the matching model, p<3x10^−4^; **Supplementary Figure 8)** and, when comparing seRNNs and L1 networks directly, seRNNs are approximated by homophily generative models better with a large effect size (*p* = 1.02x10^−40^, Cohen’s *d* = 0.80). In **Figure 2e** we further show direct comparisons data to macroscopic findings in children’s structural connectomes (left; Akarca et al., 2021) and high-density multielectrode arrays of rodent cortical neuronal networks at micrometer scales (right; Akarca et al., 2022). In **Supplementary Figure 9**, we demonstrate how homophily models increasingly approximate topology with time and regularization constraints.

To summarize so far: by incorporating spatial and communication constraints within a neural network, we produce structural topologies very reminiscent of empirical brain network architectures: modularity, small-worldness and homophilic generative properties.

### Functionally related units spatially organize in seRNNs

So far, we have explored how imposing biophysical constraints within seRNNs produces structures that mimic observed networks. However, this says nothing about the functional role of neurons or their patterning within the network. We next examined this by exploring the configuration of functionally related neurons in 3D space (**Figure 3a**). In brain networks, neurons sharing a tuning profile to a particular stimulus tend to spatially group within a spatial structure (Kanwisher, 2010; Thompson & Fransson, 2017). This occurs across spatial and functional scales, such as fine-grained cortical surfaces with preferences for stimuli features (Waskom & Wagner, 2017) (**Figure 3b**) to whole-brain functional connectivity which form regular modular network patterns (Ji et al., 2019) (**Figure 3c**). Additionally, large-scale high-resolution recordings in rodents show how the brain keeps many variable codes localized but also distributes some across the network (Steinmetz et al., 2019).

**Figure 3.**
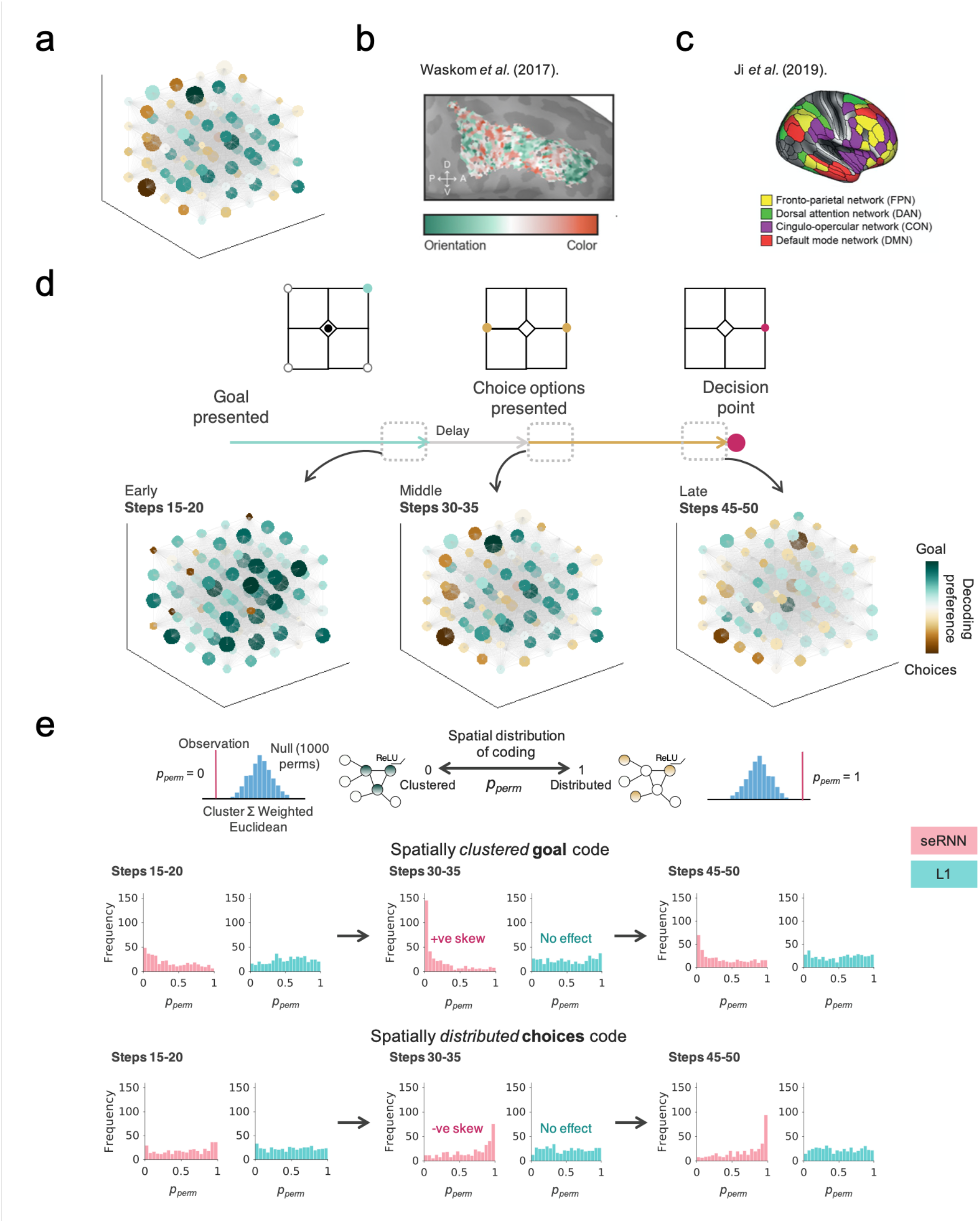
Functional clustering and distribution of coding in space. **a** An example of a representative seRNN network. The color of the nodes relates to the decoding preference of that neuron, where a preference for goal information are represented by green and choices information by brown. **b** Waskom, *et al*. (2017) highlights the spatial clustering of neuronal ensembles which are preferentially tuned for orientation (green) versus color (orange) in dorsolateral prefrontal cortex. **c** Ji, *et al*. (2019) highlights the macroscopic spatial organization of functional networks. **d** We show decoding of neuronal units for goal (green) versus choice (brown) information at different points in the trial, within the representative seRNN network. **e** A schematic illustration of the spatial permutation test for determining functional clustering (top-left) or distribution (top-right). The *p*_*perm*_ values across RNNs are given for goal information (middle) and choice information (bottom) for seRNNs (pink) and L1 networks (blue). Goal information is shown to be clustered, as given by the positively skewed *p*_*perm*_ distributions. Choice information is shown to be distributed, as given by the negatively skewed *p*_*perm*_ distributions.

To test if seRNNs recapitulate functional co-localization, we decoded how much variance of unit activity can be explained by the goal location or available choice options, over the course of each trial (see **Methods; *Decoding***). In **Figure 3d** we show a visualization in a representative network and unit-specific preferences over the course of a single trial.

By taking the relative preference for goal versus choice (calculated simply as the goal explained variance minus the choice explained variance) for each unit, we tested whether sensitivity to task-relevant information was concentrated in parts of the network. We did this via simple spatial permutations, testing if the Euclidean distance between highly “goal” or “choice” neurons was significantly less or more than would be expected by chance (see **Methods; *Spatial embedding permutation test***). A small *p*_*perm*_ value corresponds to the observation that functionally similar neurons tend to also be significantly clustered in 3D space and a large *p*_*perm*_ corresponds to functionally similar neurons being distributed in space (**Figure 3e, top**).

We tested for functional co-localization across three time-windows of the trial duration (the total duration of a trial was 50 steps; **Figure 1e**): (i) an early stage (goal presented, decoded from steps 15-20); (ii) a middle stage (choice-options presented, decoded from steps 30-35) and (iii) a late stage (decision point, decoded from steps 45-50). At the early stage, when only goal information given to the network, neurons code only for the goal information (seen by wide-spread dark green nodes in **Figure 3d, left**). In seRNNs, there is a slight positive skew in *p*_*perm*_ values, suggesting clustering of highly goal coding neurons (**Figure 3e, middle-left**). Subsequently, in the middle stage, when choice options are first presented, this goal information clusters within a concentrated area of space, leaving the choice information distributed (seen by clustering of green nodes and distribution of brown nodes in **Figure 3d, middle**). This is shown by a large positive skew in *p*_*perm*_ values for the goal in seRNN networks (**Figure 3e, middle-top**) and correspondingly the opposite for choice information (**Figure 3e, middle-bottom**). In the late stage, the clustering of goal information in space dissipates such that by the time a decision must be made the goal information has now spread out slightly more than before – although still retaining some clustering (**Figure 3e, middle-right**). At this point, the choice code has now become distributed (**Figure 3e, bottom-right**). This suggests that seRNNs use their highly modular structure to keep a strongly connected core with goal information which needs to be retained across the trial. It uses spatially proximal units to form this core. The presented choices information is then represented by units outside this core and dynamically integrates with the information in the core as the decision point approaches. Note that these findings are unique to seRNNs, shown by the distribution of seRNN *p*_*perm*_ values being skewed. L1 *p*_*perm*_ values remain uniform, indicative of no functional organization (no statistical effect). In **Supplementary Figure 10** we show that these findings are true when variables (goal, choices) are treated independently instead of relatively.

### Mixed selectivity and energy efficient coding

So far, we have shown that adding spatial constraints to a network as it learns gives rise to patterns of network connectivity which are highly reminiscent of observed biological networks. Nodes functionally co-localize, and the spatial embedding causes differences in how they code task-relevant information. This selectivity profile has been widely studied across species and brain regions. Studies show that neurons in task-positive brain regions tend to show a mixed selectivity profile, meaning that neurons do not only code for a single task variable but instead a mixture of them (Rigotti et al., 2013; Wallach et al., 2022; but see Hirokawa et al., 2019 and Whittington et al., 2022). A mixed selective code is assumed to support a network solving complex tasks by increasing the decodability of information from the network’s neurons (Fusi et al., 2016; Johnston et al., 2020). There are many ways to quantify selectivity profiles (e.g., see Bernardi et al., 2020 for recent implementations). One very simple method is to look at the correlation of explained variances per task variable across the population of neurons. These are expected to be uncorrelated, implying a neutrally mixed code where a neuron’s coding preference for one variable does not predict its code for another variable. In single unit recordings, correlations can be close to zero or sometimes slightly positive (Erez et al., 2022). In our analysis we next tested whether seRNNs show a similar uncorrelated code.

In the first instance we looked at the correlation of selectivities of trained networks (epoch 9) for the goal and choices variables. At the time point in the trial when the network must make a choice, the median correlation falls on r = -0.057 for seRNN but r = -0.303 for L1, showing that L1 networks tend to produce an anti-correlated code while seRNNs have a nearly purely mixed selective code (**Figure 4a**). It is possible that this effect is driven by the differential connectome structure of the two groups of networks. While a purely modular and separated network would not automatically also mix codes across variables evenly, we find a well-mixed code in seRNNs. The additional highly communicative connections between modules of the small-worldness characteristic might help seRNNs to organize units in space while retaining a mixed code across the population.

**Figure 4.**
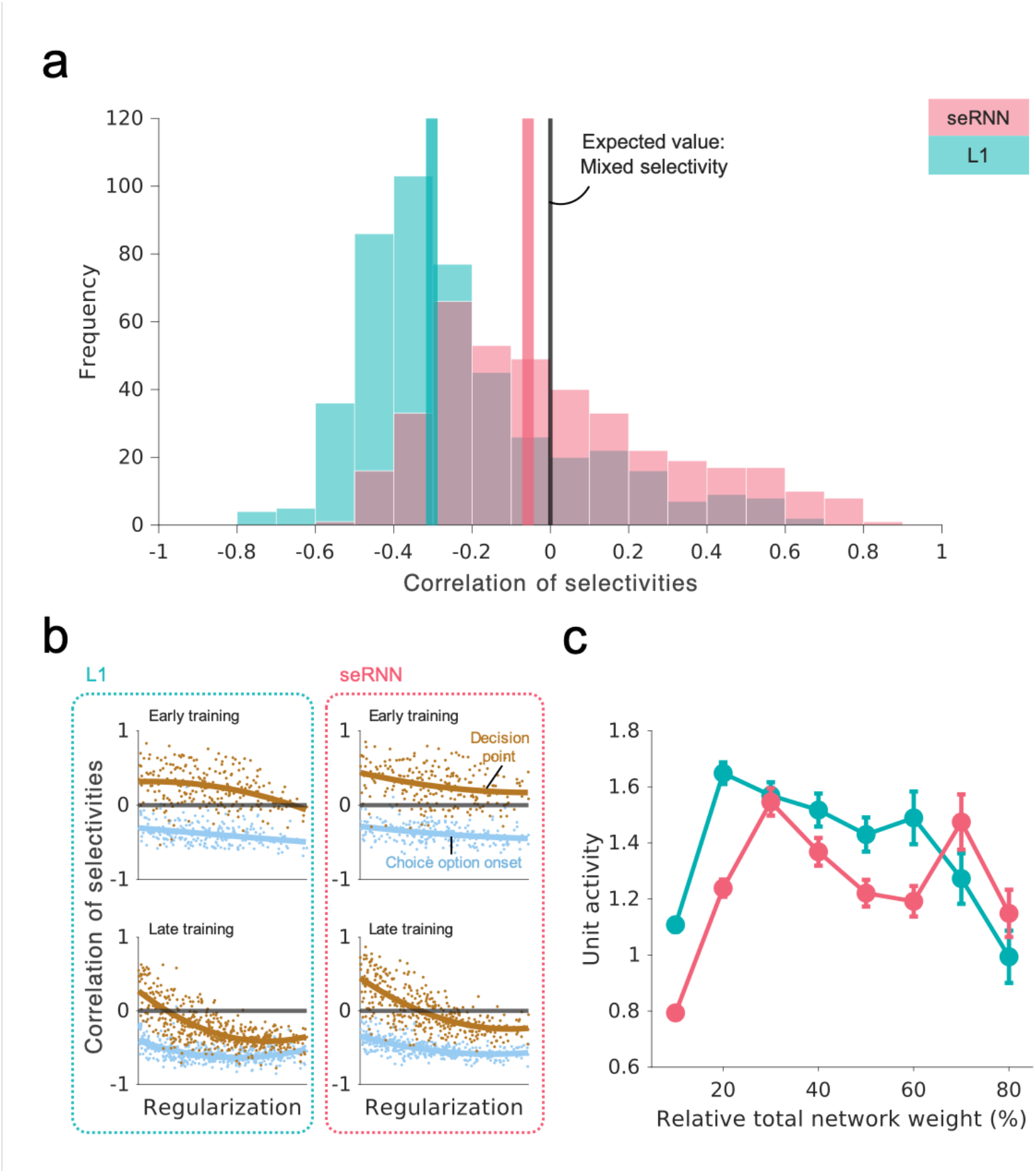
Mixed selectivity and energy efficiency. **a** The histogram of correlations of selectivities at the decision point (correlation between explained variance for goal and explained variance for choices) shows how the distribution of seRNNs is more centered around the expected value r = 0 than the L1 networks. Colored lines mark the median of the distribution. The expected value corresponds to a fully mixed selective code. Colored lines indicate the median values of the distributions. **b** Longer training and stronger regularization both cause a more anti-correlated code. At the point of decision, seRNNs naturally center on a purely mixed selective code as they do not go far below r = 0, unlike L1 networks. **c** seRNN networks spend less energy on unit activations than L1 networks which are matched for mean weight strength in the recurrent layer.

Next, we explored the correlation of selectivities across epochs, regularization strengths and time points within the trials. Looking at the correlations across the regularization spectrum, corresponding to the strength of the spatial embedding, in epoch 9 (**Figure 4b, lower row, ochre)**, stronger regularization led to an anti-correlated code in both L1 and seRNN, but seRNNs’ negative slope leveled off more quickly than L1, protecting them from becoming too anti-correlated. With regards to training time, codes became more anti-correlated with higher epoch number, as training duration is related to stronger regularization (**Figure 4b, upper row with lower row**). We also saw the correlation of selectivities changing within a trial, starting with an anti-correlated code which is becoming more mixed throughout the trial as the point of decision approaches (**Figure 4b, compare blue lines with ochre lines**). This likely is a good strategy for the network, as an initially segregated goal code makes sure that goal information does not vanish too quickly. As the trial goes on, the goal code gradually mixes with the choices code to arrive at a highly informative mixed selective code when the network needs to infer the correct choice. Like our structural results, we saw that there is significant variation across the population of networks for both L1 and seRNNs (see **Figure 4a, b**), where some networks fall neatly on r = 0 and others might show strongly correlated codes. The following section provides an analysis of which specific networks show mixed selective codes.

The choice of a neuronal code in a population of neurons is strongly linked to the question of energy demand. As the firing of action potentials uses a significant portion of energy (Attwell & Laughlin, 2001), a population of neurons should choose a code with a good trade-off of metabolic cost and information capacity (Johnston et al., 2020). We wanted to investigate whether the structural and functional differences identified so far have an impact on the amount of energy spent on signals sent through the network. To test this, we calculated the mean activation of each unit in a network’s recurrent layer (epoch 9 networks) during the period of information integration (after the onset of choices). Then we tested for the difference between seRNNs and L1 networks, controlling for the effect of the average weight strength in the recurrent layer (**Figure 4c**). seRNNs spend significantly less energy on unit activations compared to L1 networks (*p* < 0.001). Sustaining a mixed-selective code at the time of choice might help the downstream integration units to decode information more easily so that fewer unit activations are needed to effectively communicate the correct choice. Note that the effect disappears for the networks with higher average weights as these are also the networks with relatively weak regularization and hence weaker spatial embedding.

The seRNN, with its brain-like constraints, seems to automatically converge on a mixed selective code at the point of choice, as we would expect to observe in brains (Erez et al., 2022; Johnston et al., 2020). The seRNN is not only closer to the brain’s code than we observe in L1 networks, but also achieves its functionality spending less energy on unit activations. The result is that seRNNs solve the task with a code that maximizes decoding performance while minimizing energy consumption. Put simply, it is not just that neurons functionally co-localize, their individual tuning properties are also shaped to minimize costs.

### Regularization leads to convergent brain-like topology and function

So far, we have seen that seRNNs show a collection of features which are commonly observed in brains but have not been related to each other before. The caveat not addressed so far is that for any feature we observed in seRNNs we also see strong variation across the population of networks (e.g., **Figure 2c** for modularity or **Figure 4a** for mixed selectivity). This opens the possibility that these features *do not* actually arise in parallel in seRNNs but instead each feature could emerge in its unique subgroup of networks. This would be unlike biological brains which exist in a critical sweet-spot area (Beggs, 2008) where all the features described in this paper are observed. In this section we tested whether all these seRNN features co-appear in a similar subset of all trained networks, defined by a unique combination of training parameters.

To study the co-occurrence of brain features in seRNNs we looked at the distribution of feature magnitude across the space of training parameters (regularization strength, number of training epochs passed). **Figure 5a** shows matrix plots for accuracy (left), total sum of weights (middle-left), modularity (middle-right) and small-worldness (right) across the entire spectrum of training epochs (x-axis) and regularization strengths (y-axis). As before, there is substantial variation in the magnitude of features across the population of networks, but now we also see that this variation is structured.

**Figure 5.**
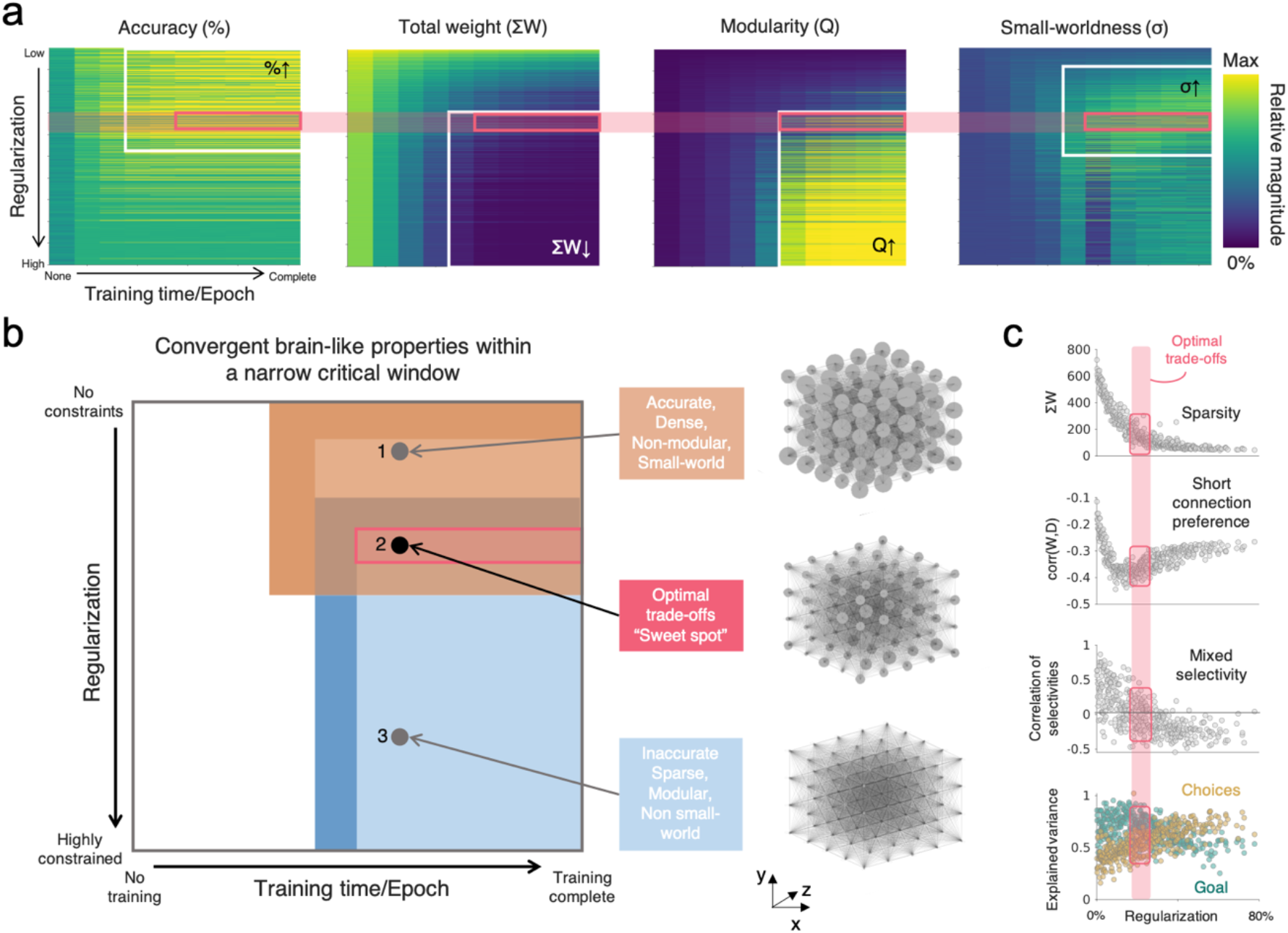
The spatially-embedded recurrent neural network (seRNN) parameter space converges on brain-like topology and function. **a** The white borders within the regularization-training parameter space delineate the conditions where seRNNs achieve robust accuracy (left), sparse connectivity (middle, left), modular networks (middle, right) and small-worldness (right). The pink box shows where all these findings can be found simultaneously. The color of the matrix corresponds to the relative magnitude of the statistic compared to the maximum. **b** This is further highlighted by a schematic representation which shows the space of possible seRNNs. The pink box shows the overlap of all findings, where accurate, sparse, modular, small-world networks are generated, which we term as being at the optimal trade-off. Network 1, 2 and 3 each represent example networks across the space. The nodes of the representative graph reflect the node’s strength, defined as the total sum of the node’s in-and out-connection weights. **c** In this pink window, networks are sparse (top), prefer short connections (middle-top), have a correlations of variable selectivities centering around zero, consistent with mixed selectivity (middle-bottom) and have equivalent explained variance for both the goal and the choice (bottom).

Brain-like topology emerges in a sweet-spot of low to medium strength regularization and during the later training epochs (pink box). The schematic in **Figure 5b** highlights this space of sparse, highly accurate, modular small-world networks with an example network exhibiting all properties (**Figure 5b, middle-right**). Above this space (i.e., networks with less regularization, highlighted in orange) networks can solve the task and exhibit small-worldness, but remain very dense and lack the modular organization found in empirical brain networks. Below this space (i.e., networks with more regularization, highlighted in light blue) networks exhibit extreme sparsity and modularity, but fail to functionally converge on the task and they lose their small-world topology. Example networks sampled from these spaces are highlighted (**Figure 5b, top-right & bottom-right**).

In a next step we wanted to look at the same “sweet spot” in terms of the network’s functional properties. The decoding required us to focus this analysis on networks with high task performance (see **Methods; *Decoding from unit activations***). We now present networks with an accuracy > 90% at epoch 9. **Figure 5c** shows the functional results across regularization strengths, highlighting the sweet-spot of regularization from **Figure 5a** with the pink box. In the first two plots from the top, we show two structural metrics (sparsity and short connection preference). We observed the same distribution when looking at the homophily generative wiring rule across the regularization spectrum (**Supplementary Figure 8b**). Looking at mixed selectivity (**Figure 5c, 3**^**rd**^ **from top**) our analyses revealed that networks show a fully mixed-selective code at the decision point in the sweet-spot window identified before. Units here show a balanced code with information for both goal and choices (**Figure 5c, bottom**), whereas very dense or very sparse network show a preference for either goal or choices information. As such the density and related modular small-world structure influences the time horizon of information flowing through the network. Here dense networks show greater focus on past information which resonates with how functional networks reconfigure to support memory (Cohen & D’Esposito, 2016). **Supplementary Figure 11** shows a detailed correlation matrix showing all pairwise relationships between features studied in the paper.

Our findings show that there is a critical parameter window in which both structural and functional brain features jointly emerge in seRNNs. Brains are often said to live in a unique but critical niche where all characteristics needed to support their function can exist in parallel (O’Byrne & Jerbi, 2022). seRNNs show the same preference for a critical parameter window but also give us the ability to study networks on their way to converging on brain-like characteristics in the critical parameter window or networks which lie outside the sweet-spot.

## DISCUSSION

Functioning brains have key organizational features endowing them with computational capacities in order to perform a broad range of cognitive operations efficiently and flexibly. These include sparse connectivity with a modular small-world structure (Bassett & Bullmore, 2017; Park & Friston, 2013; Yan & He, 2011) that are generatable via homophilic wiring rules (Akarca et al., 2021; Betzel et al., 2016; Vértes et al., 2012) with spatially-configured functional units that implement a mixed-selective code (Fusi et al., 2016; Rigotti et al., 2013) which concurrently minimizes energy expenditure (Attwell & Laughlin, 2001; Johnston et al., 2020). We argue that these complex hallmarks can be, at least in part, attributed to three forces impacting virtually any brain network: Task control, structural costs and local communication constraints. In this work we have shown that spatially-embedded recurrent neural networks (seRNNs) allow us to experimentally manipulate these constraints, demonstrating that seemingly unrelated neuroscientific findings can emerge in unison and appear to have a strong codependence.

The theoretical backdrop to this convergence of phenomena lies in conceptualizing the brain as having to resolve multiple “economic” challenges at once. That is, given limited metabolic resources, the brain must perform as well as possible given the environment it finds itself in (for related ideas on resource-rationality, see Gershman et al., 2015; Kool & Botvinick, 2018; Todd & Gigerenzer, 2012). We hypothesized that numerous structural and functional neuronal features found in brains across species could be the result of this very fundamental optimization and how it unfolds. Indeed, the predominant explanation for anatomical organization of the brain has focused on minimizing wiring costs while maximizing adaptive topological features (Bullmore & Sporns, 2012; Zhou et al., 2022). As such, we have seen considerations of space implemented in functional feedforward neural network models (Gozel & Doiron, 2022; Huang et al., 2019; Lee et al., 2020). Here we instantiated our core hypothesis mathematically within the seRNN model by providing two challenges to RNNs during supervised learning: (1) long connections should be minimized where possible – reflective of their metabolic cost (Kaiser & Hilgetag, 2006; Sporns, 2011), and (2) connections can only change their weights as a function of their underlying communication – reflective of signal propagation between neuronal units (Betzel et al., 2022; Seguin, Jedynak, et al., 2022; Seguin, Mansour L, et al., 2022; Shimono & Hatano, 2018). Both challenges are addressed at the local neuronal-level over the course of training which has the effect of continually shaping the networks global structural and functional properties over time.

Our findings show that, within a critical window of regularization, seRNNs recapitulate numerous empirical structural and functional neuroscience observations simultaneously. This suggests that providing artificial neural networks with a topophysical structure (Bassett et al., 2010; Sperry et al., 2017) can enhance our ability to directly link computational models of neural structure and function. Notably, recent work placing empirical connectomes within a reservoir framework made important steps towards this goal by partially fixing the structure of the network (Damicelli et al., 2021; Goulas et al., 2021; Suárez et al., 2021). In contrast, our seRNNs approach allows us to be very specific in defining the networks’ constraints and objectives, in both physical structure and function, while allowing the network’s connectivity to change over time and hence simulate more realistic interactions between network structure and function.

Our findings extend beyond neuroscience and have implications for modern artificial intelligence (AI) research. A technical summary of our approach is that we have introduced a regularization function pushing RNNs to converge on brain-like sparse structure and function, while they are trained to optimize task function via a regular backpropagation algorithm. Importantly, sparse models also play a large role in machine learning and AI. One of the first widely used implementations of sparsity can be found in regression models, specifically LASSO regression (Tibshirani, 1996). Regularization can generally be interpreted as a prior over the space of possible parameter values (Hardt & Recht, 2022) and in the case of LASSO regression, the L1 regularization is nudging the model to converge on a generally sparse model. As modern neural network models increasingly have grown in size and complexity (Ramesh et al., 2022; Reed et al., 2022), sparsity has received a new wave of attention because a reduced set of parameters can make training and storing a model more efficient (Hoefler et al., 2021) and allow for processing of longer input sequences (Zaheer et al., 2021). We have seen this development especially in vision models (Han et al., 2015; Zhang et al., 2022) and also recently in reinforcement learning (Graesser et al., 2022). Generally, these studies show that large neural networks can easily be pruned without negatively impacting task accuracy. Our findings expand this literature by demonstrating that neuroscience can inform the specific shape and structure of the trained model (Chechik et al., 1998; Lindsay et al., 2017). For example, our results and related simulations from other groups show how the network’s structure influences the selectivity profile of a network’s units (see **Figure 5**; Lindsay et al., 2017). seRNNs converge on a spatially structured code but preserve neural diversity of signals by keeping the signals of cells across the population mixed-selective. Regular neural networks, like our L1 baseline models, can lack richness in the final learned representations (Papyan et al., 2020) which recently has been hypothesised to inhibit a model’s performance in solving complex problems (Ma et al., 2022). As such, seRNNs highlight how neuroscience can provide a lens by which to observe how concepts of biological structure and function can aid efforts to overcome limitations in neural networks.

There are many areas that we wish to improve upon with future research. Principally, our models did not include a significant amount of biological detail that, while inevitably critical for neuronal functioning, do not speak to the specific observations we aimed to recapitulate in the present study. Nevertheless, implementing such details including specific molecular mechanisms that guide neural circuit development (Moons & De Groef, 2021) or heterogeneous spiking of neurons (Perez-Nieves et al., 2021) will likely provide a range of new insights into the trade-offs specific to biological brains. Indeed, the addition of such biological details will help us expand the applicability of our models to explore the effect of developmental time-courses (Baxter & Levy, 2020; Chechik et al., 1998), functional brain specialization (Johnson, 2011) and how network variability may underpin individual differences (Siugzdaite et al., 2020). Beyond biological detail, there are also numerous computational directions that are yet to be examined – for example, currently we do not allow units in the seRNNs to reposition themselves in space, nor do we train networks to make continuous choices in a multi-task environment.

The development of spatially-embedded RNNs allowed us to observe the impact of task control, structural cost and communication constraints in a model system which is able to dynamically trade-off its structural and functional objectives. This showed us how a wide selection of neuroscientific findings, which have not been linked to each other before, arise in unison during the same optimization process. These features are: Sparse connectivity, preference for short distance connections, modularity, small-worldness, homophily, the configuration of functionally similar units in space, mixed selectivity and an energy efficient code. We believe that the modelling approach shown to work in seRNNs will be able to speed up innovations in neuroscience by allowing us to systematically study the relationships between features which all have been individually discussed to be of high importance to the brain.

## METHODS

### Task paradigm

The task that networks are presented with is a one choice inference task. Networks first observe stimulus A for 20 time-steps, followed by a delay for 10 time-steps, followed by stimulus B for 20 steps. Agents must then make one choice. This setup can be interpreted as a one-step navigation task, where agents are presented with the goal location (stimulus A) followed by possible choice directions (stimulus B). The choice to be made is the one moving closer to the goal. The following table outlines all possible trials and defines whether the given trial is included in the baseline version of the task, compared with the hard version (see **Supplementary Figure 7**). For the randomized version of the task we randomly shuffle the correct choice across the entire training set of trials so that networks cannot learn to make correct choices above chance.

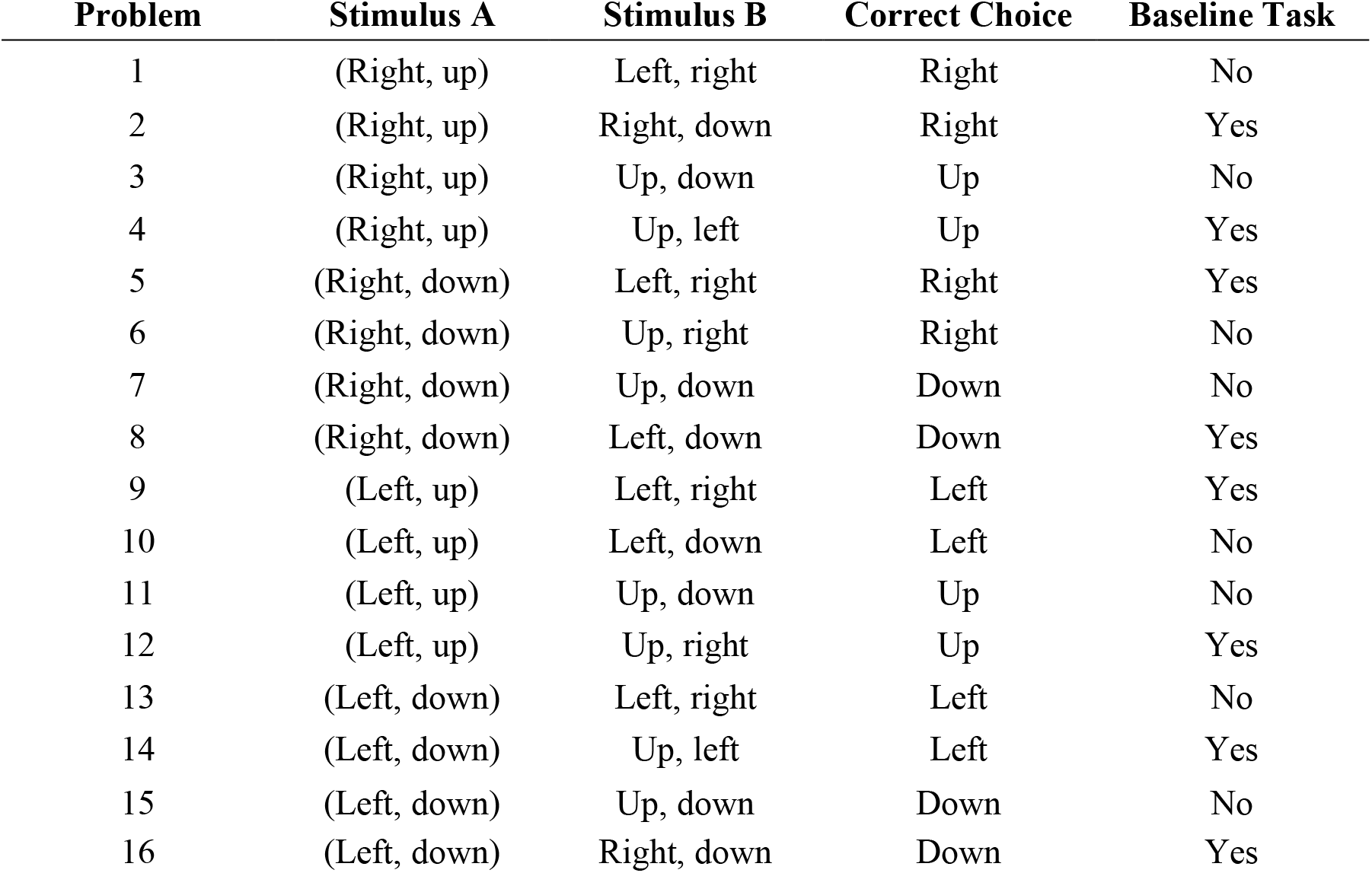

| Problem | Stimulus A | Stimulus B | Correct Choice | Baseline Task |
| --- | --- | --- | --- | --- |
| 1 | (Right, up) | Left, right | Right | No |
| 2 | (Right, up) | Right, down | Right | Yes |
| 3 | (Right, up) | Up, down | Up | No |
| 4 | (Right, up) | Up, left | Up | Yes |
| 5 | (Right, down) | Left, right | Right | Yes |
| 6 | (Right, down) | Up, right | Right | No |
| 7 | (Right, down) | Up, down | Down | No |
| 8 | (Right, down) | Left, down | Down | Yes |
| 9 | (Left, up) | Left, right | Left | Yes |
| 10 | (Left, up) | Left, down | Left | No |
| 11 | (Left, up) | Up, down | Up | No |
| 12 | (Left, up) | Up, right | Up | Yes |
| 13 | (Left, down) | Left, right | Left | No |
| 14 | (Left, down) | Up, left | Left | Yes |
| 15 | (Left, down) | Up, down | Down | No |
| 16 | (Left, down) | Right, down | Down | Yes |

All stimuli are One-Hot encoded with a vector of 8 binary digits. The first 4 define the goal locations and only one of the 4 digits would be set to one during the goal presentation. The second 4 binary digits each stand in for one allowed choice direction and two choice directions would be set to one during the choice options presentation. Gaussian noise with a standard deviation of 0.05 is added to all inputs.

This task design is a simplified version of a multi-step maze navigation task we have recorded in macaques. The hard version of the task with an extended set of trials is equivalent to the first choice monkeys face in their version of the task. The monkeys then continue with a further step to reach the goal and collect the reward. As the goal of this study was to establish the emerging features of seRNNs, here we focus just on the first choice and leave questions relating to the multi-step task to future investigations.

### Recurrent neural network modelling

All recurrent neural networks in this project have 100 units in the hidden layer and are defined by the same basic set of equations:

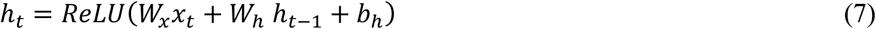

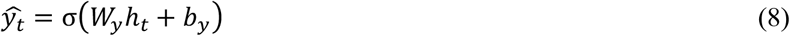

- *x*_*t*_: Input vector at time t (1x8)
- *W*_*x*_: Input layer weight matrix (8x100)
  - Xavier initialization
- *h*_*t* − 1_: Activation of hidden layer at time t-1 (1x100)
  - Zeros initialization
- *W*_*h*_: Hidden layer weight matrix (100x100)
  - Orthogonal initialization
- *b*_*h*_: Bias of hidden layer (1x100)
  - Zeros initialization
- *h*_*t*_: Activation of hidden layer at time t (1x100)
- *W*_*y*_: Output layer weight matrix (100x8)
  - Xavier initialization
- *b*_*y*_: Bias of network output (1x8)
  - Zeros initialization
- *σ*: Softmax activation function
- 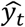: Network output / prediction

Networks differ in terms of which regularization was applied to its hidden layer (see **Results; *How to spatially embed a recurrent neural network***) and with which regularization strength. Networks are optimized to minimize a Cross Entropy Loss on task performance combined with the regularization penalty using the Adam optimizer (hyper parameters: learning rate = 0.001, beta_1 = 0.9, beta_2 = 0.999, epsilon = 1e-07) for 10 epochs. Note that the network’s choice is only read out once, at the very end of the trial. Each epoch consists of 5120 problems, batched in blocks of 128 problems.

Networks trained on the randomized version of the task (see **Supplementary Figure 7**) are trained on epochs consisting of 3840 problems to avoid them becoming too sparse too quickly which would prohibit us from analyzing the trajectory of metrics.

### Regularization strength setup and network selection

The most critical parameter choice in our analyses is the regularization strength. As shown across analyses (e.g., **Figure 2** and **Figure 5**), the strength of the regularization has a major influence on all metrics analyzed here. While the L1 regularization and the purely Euclidean regularization could be matched by average strength of regularization of the hidden layer, the communicability term of seRNNs makes this challenging due to it being dependent on the current state of the hidden layer and hence changing throughout training. To match the spectrum of regularization strengths in L1 and seRNNs we used a functional approach. As performance in the task starts to break down as networks become too sparse to effectively remember past stimuli, we matched regularization strength using task performance before looking at any of the other structural or functional metrics. Specifically, we set the regularization spectrum on a linear scale and chose the boundary values so that task performance started to significantly deteriorate half-way through the set of networks (so around the 500^th^ network for the sets of 1000 networks).

To make both groups comparable, we focus our analyses on networks which achieve >90% task accuracy. For the L1 networks these were 47.3% of all trained networks and for seRNN networks 39.2%. Note that this difference in percentages is not meaningful *per se* and could be eliminated by matching the regularization spectra of both groups more closely. As we focus our analyses on highly functional networks with high task accuracy, matching the regularization spectra of both groups would have not influenced the results. The code repository has an overview file with regularization strengths chosen for different network types. We hope that future implementations of the seRNNs can provide a method for more precise numerical matching between regularization strengths.

### Topological analysis

Graph theory network statistics were calculated using the Brain Connectivity Toolbox (Rubinov & Sporns, 2010), and the mathematical formalisms are provided. All network statistics were calculated on the hidden RNN weight matrix and all edges were enforced to be the absolute value of the element. When the measure in question was binary (e.g., small-worldness) a proportional threshold was applied, taking the top 10% of these absolute connections.

#### Modularity

The modularity statistic, Q, quantifies the extent to which the network can be subdivided into clearly delineated groups:

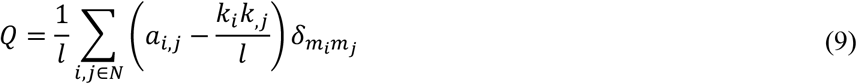

where *m*_*i*_ is the module containing node *i*, and 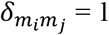 if *m*_*i*_ = *m*_*j*_, and 0 otherwise. In this work, we tested the modularity using the default resolution parameter of 1.

#### Small-worldness

Small-worldness refers to a graph property where most nodes are not neighbors of one another, but the neighbors of nodes are likely to be neighbors of each other. This means that most nodes can be reached from every other node in a small number of steps. It is given by:

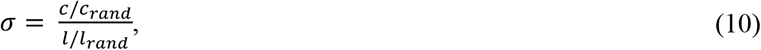

where *c* and *c*_*rand*_ are the clustering coefficients, and *l* and *l*_*rand*_ are the characteristic path lengths of the respective tested network and a random network with the same size and density of the empirical network. Networks are generally considered as small-world networks at σ > 1. In our work, we computed the random network as the mean statistic across a distribution of n = 1000 random networks. The characteristic path length is given by:

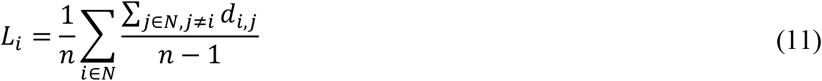

#### Degree

The degree is the number of edges connected to a node. The degree of node *i* is given by:

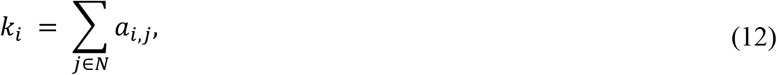

where *a*_*i, j*_ is the connection status between *i* and *j. a*_*i, j*_ =1 when link *i*, exists (when *i* and *j* are neighbors); *a*_*i, j*_ = 0 otherwise (*a*_*i, i*_ = 0 for all *i*).

#### Clustering coefficient

The clustering coefficient is the fraction of a node’s neighbors that are neighbors of each other. The clustering coefficient for node *i* is given by:

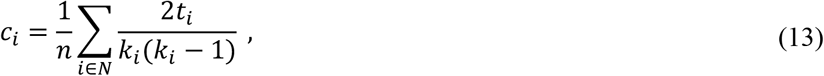

where *c*_*i*_ is the clustering coefficient of node *i* (*c*_*i*_ = 0 for *k*_*i*_ < 2).

#### Betweenness centrality

The betweenness centrality is the fraction of all shortest paths in the network that contain a given node. Nodes with high values of betweenness centrality therefore participate in a large number of shortest paths. The betweenness centrality for node *i* is given by:

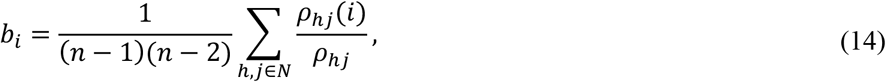

where *ρ*_*hj*_ is the number of shortest paths between *h* and *j*, and *ρ*_*hj*_(*i*) is the number of shortest paths between *h* and *j* that pass through *i*.

#### Edge length

The edge length is the total edge lengths connected to a node. It is given by:

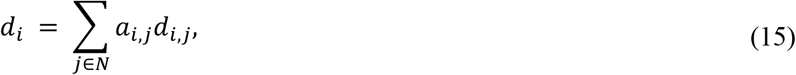

where *d*_*i, j*_ is the Euclidean distance between *i* and *j*.

#### Global efficiency

The global efficiency is the average of inverse shortest path length. It is given by:

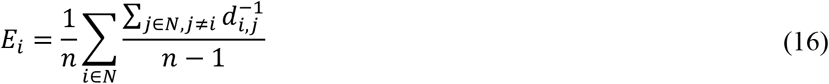

#### Matching

The matching index computes the proportion of overlap in the connectivity between two nodes. It is given by:

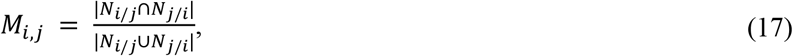

where *N*_*i*/*j*_ refers to neighbors of the node *i* excluding node *j*. Where global measures of matching have been used, we averaged across the upper triangle of the computed matching matrix.

### Generative network modelling

To elucidate if the topology of our RNNs could be recapitulated via unsupervised wiring rules analogous to published works (Akarca et al., 2021; Betzel et al., 2016; Vértes et al., 2012) we attempted to simulate their topology via a generative network model.

The generative network model (GNM) can be expressed as a simple wiring equation (Akarca et al., 2021; Betzel et al., 2016; Kaiser & Hilgetag, 2004; Vértes et al., 2012) in which connections are probabilistically added to the network in discrete time-steps. Starting from an empty network, at each time point a connection is added probabilistically according to the following simple topological wiring equation:

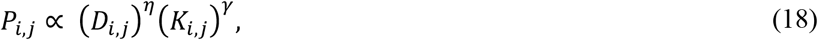

where *D*_*i, j*_ represents the Euclidean distance between nodes *i* and *j* (as outlined above), and *K*_*i,j*_ reflects some topological value in forming a connection. We tested 13 established *K*_*i,j*_ wiring rules that have been studied elsewhere extensively (Akarca et al., 2021, 2022; Betzel et al., 2016; Carozza et al., 2022; Oldham et al., 2022) **(Table 1)**. *P*_*i, j*_ represents the wiring probability as a function of the product of the parameterized costs and topological value. The end-result is a wiring probability matrix which updates over time as new connections are added. Wiring parameters *η* and *γ* are each scalars which tune the relative influence of costs and value on the wiring probability.

**Table 1.**
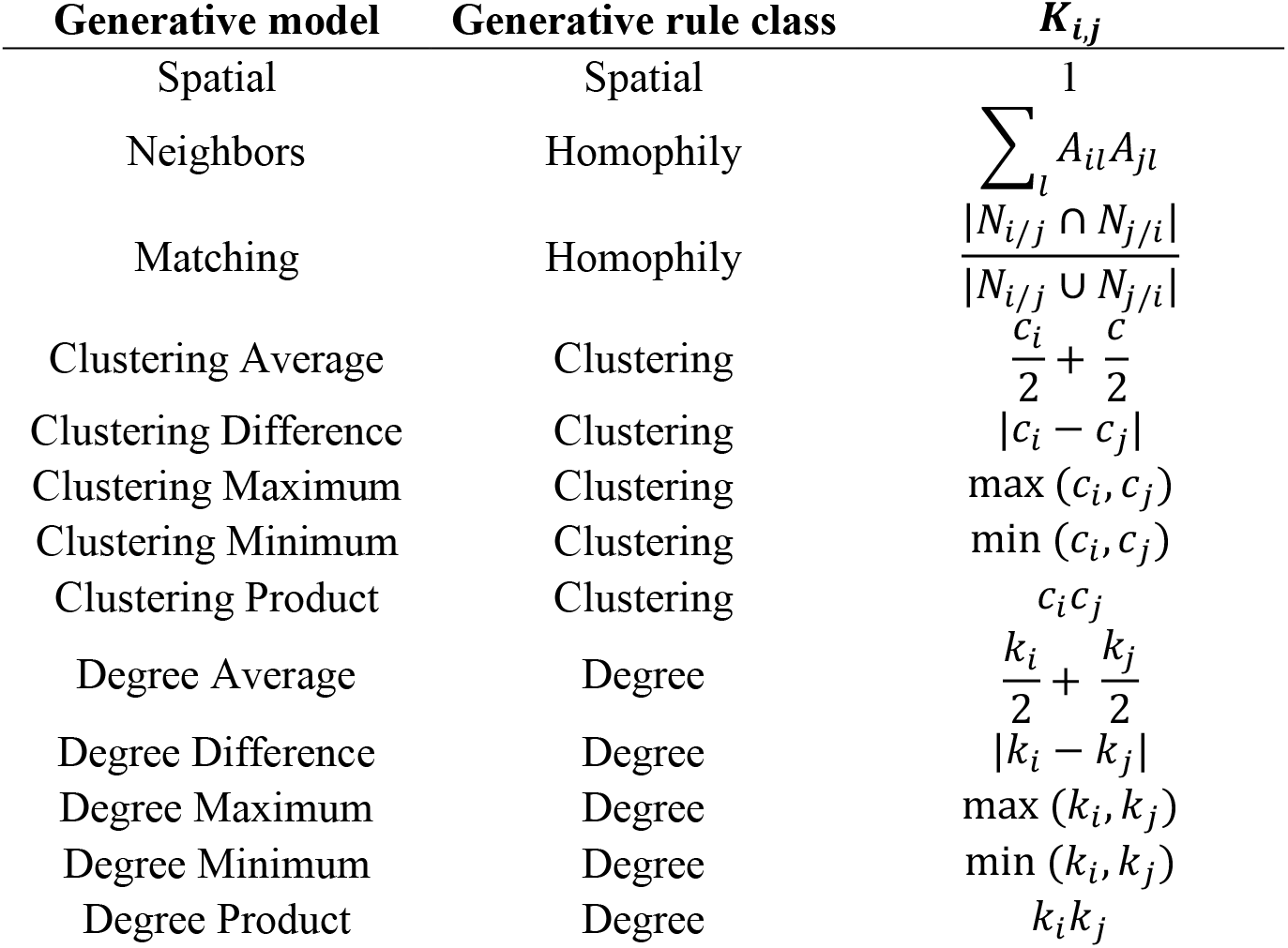
Topological wiring rules used in the generative network model.

### Voronoi tessellation parameter fitting procedure

To generative synthetic networks, we ran simulations across a defined a parameter space of 1000 evenly spaced combinations (-3 ≤ *η* ≤ 3 and -3 ≤ *γ* ≤ 3), for each of the 13 generative wiring rules (**Table 1**). This produced a total of 13,000 simulations that were compared to each of the RNNs. Of the 1000 RNNs for L1 and seRNNs (sorted by marginally increasing regularization), we constrained our analysis between the 100^th^ network (which had been relatively weakly-regularized) to the 600^th^ network (which had been relatively strongly-regularized) that were functional (as defined as having >90% validation accuracy), which left 455 and 379 networks for L1 and seRNN networks respectively. This left 834 RNNs for each of the sets, each with 13,000 simulations, leading to a total of 10,842,000 evaluated simulations in total. It is important to note that the generative modelling process is based on binary connections rather than weighted. As with the topological analysis, we enforced a 10% proportional threshold on the absolute weights of the RNN hidden layer to attain these binary networks.

The parameter fitting for each network was done using a Voronoi tessellation procedure (Betzel et al., 2016). This procedure works by first randomly sampling the parameter space and evaluating the model fits of the resulting simulated networks, via the energy equation. Following an initial search of 200 parameters in this space, we performed a Voronoi tessellation, which establishes 2D cells of the space. We then preferentially sampled from cells with better model fits according to the energy equation (Betzel et al., 2016). Preference was computed with a severity of α = 2 which determines the extent to which cell performance led to preferential sampling in the next step. This procedure was repeated a further four times, leading to a total of 1000 simulations being run for each considered network across all generative rules (as stated above).

### Generative model cost functions

To evaluate the fitness of synthetic networks relative to our RNNs, we used two different cost functions which each compute a measure of dissimilarity between the synthetic network and the RNN. The first cost function, termed the energy equation, computes the Kolmogorov-Smirnov (KS) distance between the observed and simulated distributions of a range of network statistics (Betzel et al., 2016). It then takes the maximum of the four KS statistics considered so that, for any one simulation, no KS statistic is greater than the energy. This is acts as a measure of dissimilarity in terms of the *global statistics* individually:

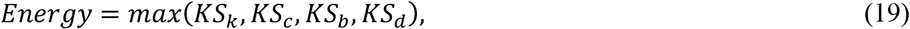

where *KS* is the Kolmogorov-Smirnov statistic comparing degree *k*, clustering coefficient *c*, betweenness centrality *b* and edge length *e* distributions of simulated and observed networks.

### Decoding from unit activations

To analyze the internal function of our trained recurrent neural networks, we record the hidden state activity of every unit while the network solves a set of 640 trials. Each trial is constituted of 50 steps (as shown in **Figure 1e**). For decoding, the activity is averaged in step-windows of 5, so that there is a total of 10 time-windows. In animal electrophysiology researchers often look at the explained variance per task variable per unit. To allow for comparison of our networks to findings in the literature, we wanted to extract the same metric. Given the nature of our task (Goal, Choice Options, Correct Choice), variables in our recordings are highly correlated, so that the standard decoding with ANOVA would give biased results. Instead, we used a decoding algorithm based on L1 regression, as follows:

1. Apply cross validated L1 regression with 5 K-Fold to set alpha term with best cross validation performance.
2. Split the dataset via repeated k-fold (3 folds, 2 repeats).
3. On each (train, test) dataset:
  1. Train L1 regression with the pre-set alpha term.
  2. Calculate explained variance in test dataset.
  3. Iteratively set all values of a given set of predictors (e.g., all goal predictors) to 0 and recalculate the explained variance and calculate the drop of explained variance per predictor group.
  4. Take mean of drop of explained variance for each group across splits of dataset.

This algorithm results in every unit in every network being assigned an explained variance number for every task variable. Note that the decoding cannot reliably work in networks which make too many errors, so that we only functionally analyze networks with a task performance of 90% or above.

### Spatial permutation testing of task-relevant information

To examine the spatial clustering of decoded task information of neuronal ensembles within the RNNs, we constructed a simple spatial permutation test as follows:

1. Considering a single RNN hidden layer at a particular task time window (note, explained variances change over the course of the task), for each neuron, compute the relative preference for goal versus choice explained variance for each unit. This is calculated as the goal explained variance minus the choice explained variance.
2. Between all *n* “goal” neurons (i.e., positive difference from step 1), compute the Euclidean distance weighted by the decoding for goal information. This therefore captures the spatial proximity between goal neurons weighted by the magnitude of their “goal” information. Average this matrix to compute a summary statistic. This is the observed statistic.
3. Then repeat this procedure for 1000 times, but for a random set of *n* neurons taken from the 3D grid space. These 1000 summary statistics are the null distribution.
4. Compute a permuted p-value (*p*_*perm*_) which is simply the location in which the observed statistic (step 2) sits within the null distribution (step 3) normalized to the range [0 1]. This value subsequently corresponds to how clustered or distributed the observed goal decoding information is clustered in space relative to random chance. A small *p*_*perm*_ means that information is clustered more than chance and vice versa.
5. Do step 1-4, but between all “choices” neurons (i.e., negative difference from step 1)
6. Redo step 1-5 for all desired time windows that have been decoded. In the current work, we calculated *p*_*perm*_ values for time window 3, time window 6 and time window 9 to reflect different aspects of the task over the sequence of the task.

The above steps were done for all functional RNNs (>90% accuracy) for L1 and seRNNs. We presented distributions of these *p*_*perm*_ values for goals and choices to highlight how goal and choices information is clustered, distributed or random at key points in the sequence of the task. To ensure that we did not bias our findings, we further computed a slight variation of the above statistical test which allows us to assess the clustering of coding information independently (i.e., without computing relative goal versus choice coding, as in step 1 above). As cluster size was now not determined by the direction of coding (as it was previously) we instead only considered nodes which coded for the top 50% of the variable (equaling 50 nodes). This was selected because this approximately mirrors the cluster sizes achieved in the primary functional clustering analysis. Mirroring the permutation testing approach, we calculated the *p*_*perm*_ by ranking the mean Euclidean distance between these nodes (top 50% coding neurons) in a null distribution of Euclidean distance between 1000 permuted samples of 50 nodes. This was done for goal, choice (to assess replication), but also the correct choice variable (which was not tested primarily, due to it being a relative measure between two variables). This test is advantageous in that it allows for testing variables independently, but disadvantageous in that it does not directly incorporate the coding magnitude into the test statistics. These findings are given in **Supplementary Figure 9**.

## Acknowledgements

We thank Matt Botvinick for helpful comments and input throughout the development of this project. J.A., Da.A., Du.A., and J.D. are supported by UKRI MRC funding. J.A. receives a Gates Cambridge Scholarship. Da.A. receives a Cambridge Trust Vice Chancellor’s Scholarship. Da.A and Du. A are both supported by the James S. McDonnell Foundation Opportunity Award. D.S. is funded by DeepMind.

## Code and data availability

With the release of the project as a preprint we provide Jupyter notebooks which show how we train our networks to perform the inference task and analyze its structure. This notebook contains the classes we use to train our networks in Tensorflow and functions for the fundamental structural analysis. This will allow researchers to run their own simulations using seRNNs. We will also provide a PyTorch implementation soon after. Upon full publication we will also make all final analysis scripts available. The project’s GitHub repository is: https://github.com/8erberg/spatially-embedded-RNN.

For the purpose of open access, the authors have applied a Creative Commons Attribution (CC BY) license to any Author Accepted Manuscript version arising from this submission.

