## Supplementary Material for "Spatially-embedded recurrent neural networks reveal widespread links between structural and functional neuroscience findings"

### CONTENTS


#### *The logic of communicability in network optimization, 3*

***Supplementary Figure 1.*** An example matrix of representation of a simple graph.

***Supplementary Figure 2.*** Computing the possible routing paths between two nodes.

***Supplementary Figure 3.*** Computing the network communicability as a function of the weighted sum of all possible paths.

***Supplementary Figure 4.*** Weighted communicability and its normalization.

***Supplementary Figure 5.*** Comparing communicability terms.

***Supplementary Figure 6.*** Changing individual weights in network to optimize the total weighted network communicability value

#### *Replication across baseline architectures, 9*

***Supplementary Figure 7.*** Independent replication of key results across a range of baseline regularizers, including various regularization and task comparisons.

#### *Generative network modelling of RNN architectures, 11*

***Supplementary Figure 8.*** L1 network generative model energy profile.

***Supplementary Figure 9.*** Homophily model fits improve with time and constraints.

#### *Spatial configuration of task-relevant variables, 13*

***Supplementary Figure 10.*** Replication of spatial configurations using a weighted variation of the spatial permutation statistical procedure.

#### *Structure-function relationships in seRNNs, 14*

***Supplementary Figure 11.*** Intertwined global structural-functional relationships in seRNNs.

#### ***The logic of communicability in network optimization***

According to graph theory, brain networks can be described as graphs that are composed of nodes (vertices) denoting a neural element (e.g., neuronal unit in an RNN or neuron) that are linked by edges representing their connections (e.g., connection in a RNN or synapse). Beyond the graph representation, it can equally be represented as a simple adjacency matrix, $A$ (**Supplementary Figure 1**).

In this example, we show a symmetrical binary graph because there are no directions in the connections and the edge has no weight – note, this is the same schematic shown in **Figure 1**.

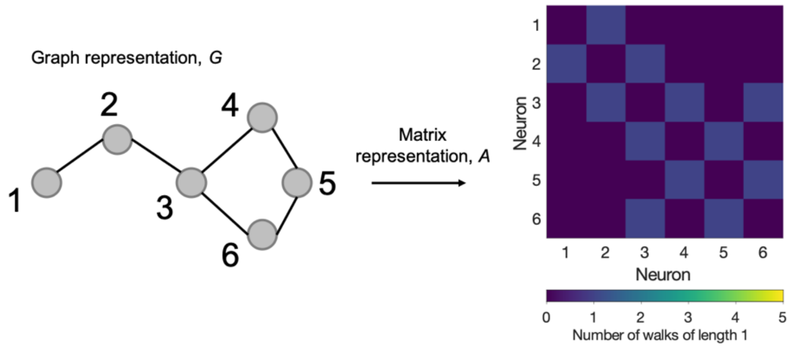


***Supplementary Figure 1.*** **An example matrix representation of a simple graph.**

There is an interesting property of graphs like this, where if you raise $A$ to the power of $k$ (i.e., $A^k$) the result tells you how many walks of length $k$ are possible between any two nodes (by walks we mean simply how many "jumps" can be taken along the graph from one node to the next). For example: $A^2$ tells you how many walks of length *2* exist between any two nodes, and $A^3$ tells you how many walks of length *3* exist between any two nodes (**Supplementary Figure 2**).

You can check this by taking any two nodes (e.g., node 5 and node 3) and count how many times you can walk a number of paths (e.g., 2 paths). In this example the answer is 2: the path can go from 5→4→3 or 5→6→3. This is reflected in the matrix $A^2_{5,3}$ which is equal to 2.

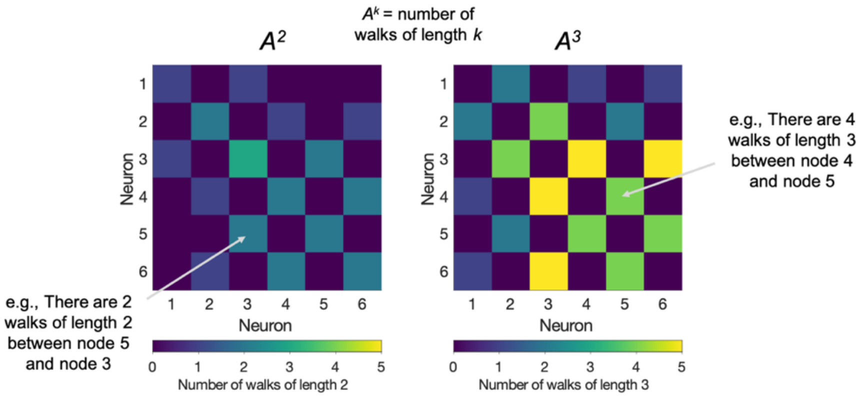


***Supplementary Figure 2*. Computing the possible routing paths between nodes.**

Why is this relevant? Because if you want to understand the extent to which two regions in a network can *communicate* with each other, you need to model the extent to which information can be passed between nodes on the graph.

To calculate this, you can simply take a *weighted sum of all possible walks between two nodes*. Note that it must be a weighted sum because as you extend the number of walks between two regions, it is less likely that any one journey was taken by that route. Given our previous example, you can now see that this would look like this:

$$\sum_{k=0}^{\infty} \frac{A^k}{k!} = \frac{A^0}{0!} + \frac{A^1}{1!} + \frac{A^2}{2!} + \frac{A^3}{3!} + \frac{A^4}{4!} + \frac{A^5}{5!} + \frac{A^6}{6!} + \frac{A^7}{7!} + \cdots$$

This expression is an infinite sum. This expression is equivalent to the Maclaurin series (Taylor series, centered at zero) of $e^A$:

$$\sum_{k=0}^{\infty} \frac{A^k}{k!} = e^A$$

An illustration of this can be seen in **Supplementary Figure 3**:

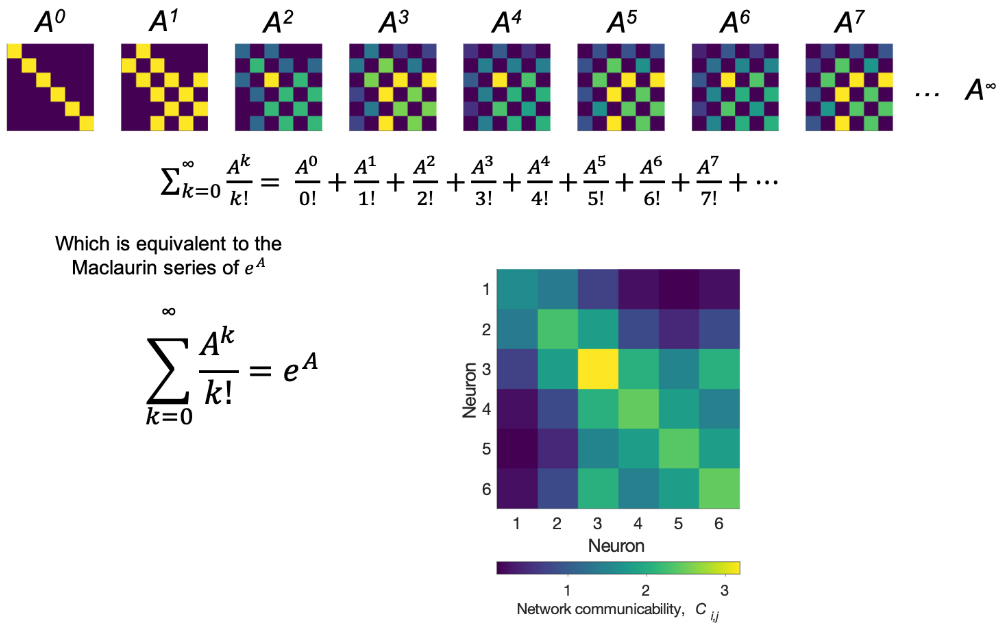


***Supplementary Figure 3*. Computing the network communicability as the weighted sum of all possible walks.**

Note that *A* here is the same original matrix we had at the beginning. So, if you raise *e* to the power of the *A* matrix, you are computing the weighted sum all possible walks between two regions – and this is what we call the *network communicability, C*. This type of communication actually resembles random diffusion or random-walks across the graph: mirroring what is a likely constraint within biological neural networks – signals propagating through the network and reaching nodes as a function of how communicable they are, based on the underlying connectivity.

The same logic can easily be applied to weighted graphs. The only difference now is that the weights also modulate the extent to which nodes communicate with each other. However, this introduces a slight problem: this equation has the property of biasing walks very heavily to single edges which may be larger – leading to undue influence on the calculation. Crofts & Higham (2009) proposed a solution where you add a normalization step which controls for each node's strength, leaving the following expression (for an example, see **Supplementary Figure 4**):

$$C = e^{S^{-\frac{1}{2}}WS^{-\frac{1}{2}}},$$

Where *S* is a diagonal matrix denoting the node's strength. This is the expression given as **Equation 6**, as a constraint to learning within the seRNN framework.

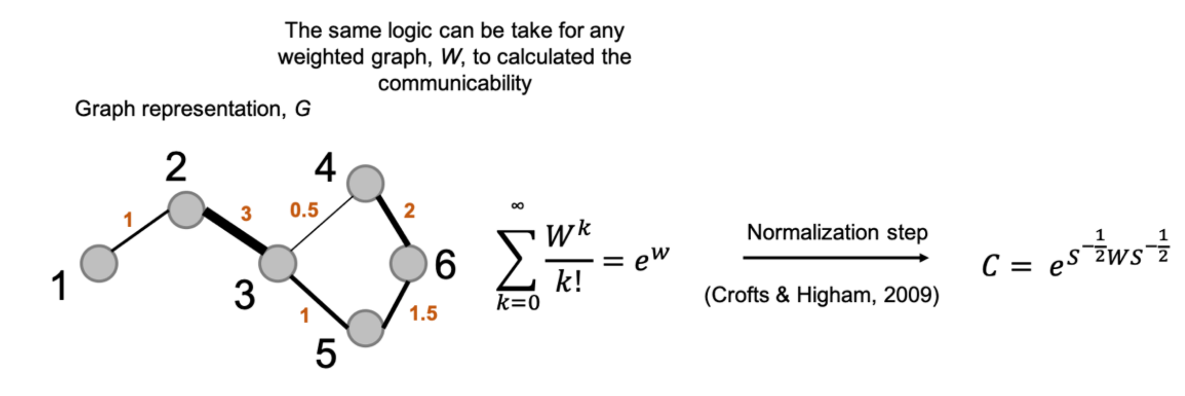


***Supplementary Figure 4.*** **Weighted communicability and its normalization.**

This normalization step has the effect of increasing the resolution of the communicability metric as shown in **Supplementary Figure 5** (for more detail, refer to Crofts and Higham, 2009). Whereas in the unweighted case highly communicative connection (e.g., between nodes 2 and 3) dominate the communicability matrix, the weighted communicability is more sensitive to individual edges' contributions to the network communicability.

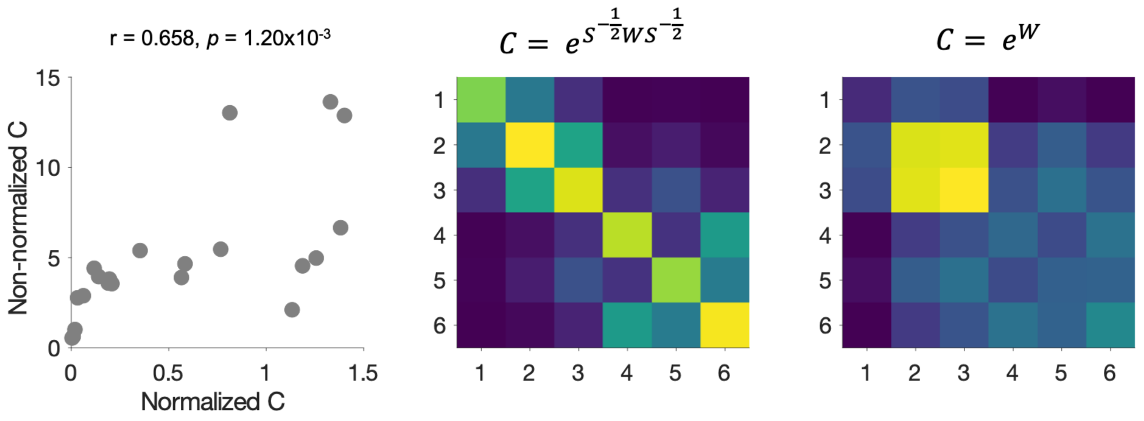


***Supplementary Figure 5*. Comparing communicability terms.**

Additionally, the weighted communicability has a desirable effect on the network optimization process as shown in show in the experiment in **Supplementary Figure 6**.

In this experiment, individual weights within the example network (top) were increased or decreased in increments of 5% of their value, and the total communicability across the network ($\sum C$) was calculated at each iteration. As shown in the scatter plot (bottom), as weights are increased (i.e., $\delta w_{ij}$ is positive – to the right) edges which have high communication due to both their high weight and position within the network (e.g., the orange edge, $w_{\{2,3\}}$) contribute towards the network becoming less communicable in total. This is because communication now gets biased towards this edge and away from other edges.

In contrast, as weights are decreased (i.e., $\delta_{ij}$ is negative – to the left), the opposite occurs; edges which have low communication (e.g., the dark blue edge, $w_{\{1,2\}}$) contribute to the network becoming less communicable in total. This means that when seRNNs are optimized for their weighted communicability (i.e., the direction of the optimization being in the down direction), highly communicable edges are strengthened and lowly communicable edges are weakened.

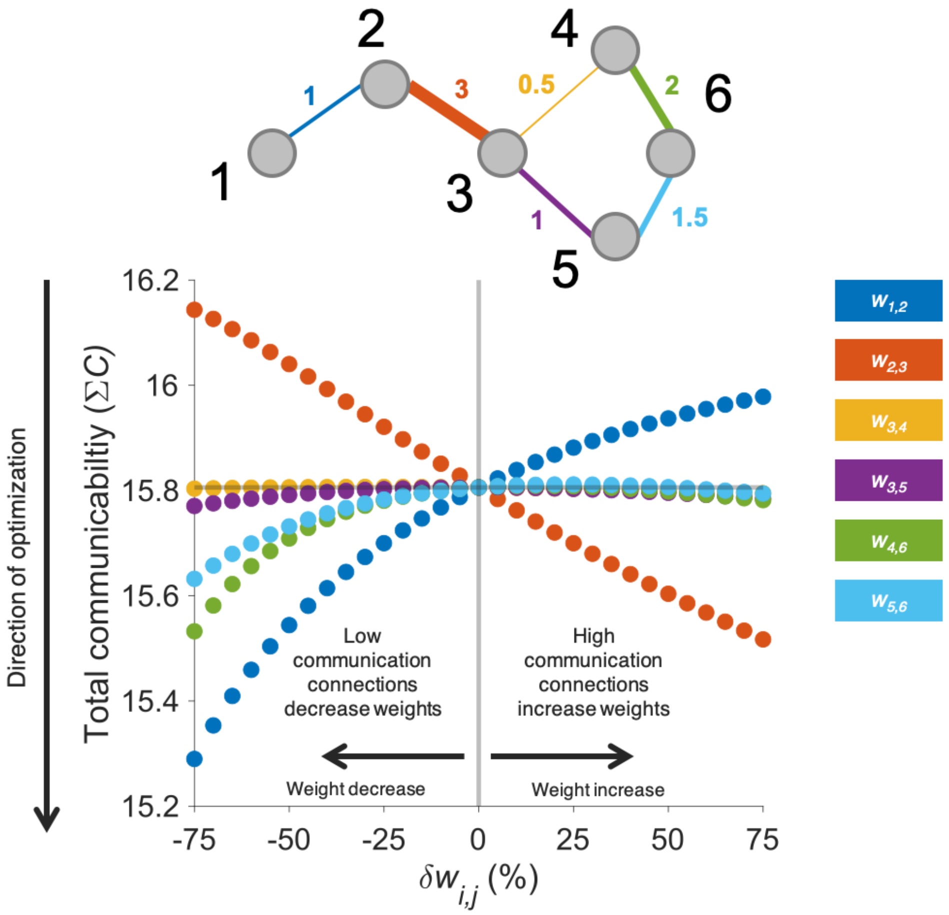


***Supplementary Figure 6.*** **Exploring the effect of changing individual network weights on the total weighted network communicability.**

#### ***Replication across baseline architectures***

Our analysis has focused on constraints we argue are ubiquitous to any biophysically instantiated neural system, modelled together and incorporated as a single optimization within a neural network architecture. But it is possible that space and communicability *have distinct roles* in outcomes that our analysis has not yet uncovered. Here we examined the effects of variations in the neural network architecture by constructing several alternative baseline regimes (see **Methods; *Baseline regimes*** for detail). First, we considered the independent effects that space ($D$only) and communication ($C$only) may have on topology (**Supplementary Figure 7a**). Broadly, we find that seRNNs show the largest magnitude in topological measures across training and L1 shows the least, in terms of total weight, weight-distance correlations, modularity and small-worldness – with the independently constrained variations falling somewhere in between. We see here that seRNNs show especially strong weight-distance anti-correlations, even though its spatial embedding is the same as for the $D$only model. We have seen that modularity is very directly related to sparsity (shown in **Figure 5**) but here we see that seRNNs are the most modular while other networks are sparser. Additionally, negative weight-distance correlations do emerge from communicability-only networks, indicating that local interactions within a physical space may also approximate some limited spatial constraints indirectly. Moreover, small-worldness occurs in absence of spatial constraints and seems to be strongly driven by the communication factor.

Beyond incorporating spatial and communication constraints, seRNNs can be conceptualized as combining both functional and structural objectives within a singular unified optimization process. As such, it is possible to ask: what is the contribution of functional goals versus physical constraints on network outcomes? To assess how the task-solving process and the task itself shapes topology, we tested two additional variations to the task in seRNN networks: (1) a harder version of the task in which there were greater variation of trials available and (2) one in which the task structure was entirely randomized, detaching goal and choice information from each other. In the comparison of functional requirements, we find that broadly there are subtle changes to the topology depending on task requirements across all considered networks (**Supplementary Figure 7b**). Harder task requirements (shown in blue) retain weights more than seRNNs, have subtly shorter connections, less modularity but retained small-worldness. The absence of a task requirement (shown in grey) leads to massively sparse networks, subtly less preference for short connections and, over training, become less small-world in their topology.

In the main text we saw that structural and functional features emerge in unison in seRNNs in a sweet spot of the strength of the spatial embedding. Here we showed that this shared emergence is also critically dependent on the process of learning the task. Additionally, we observe that the characteristic features of biological networks are caused by the joint influence of structural constraints and functional objectives on the network.

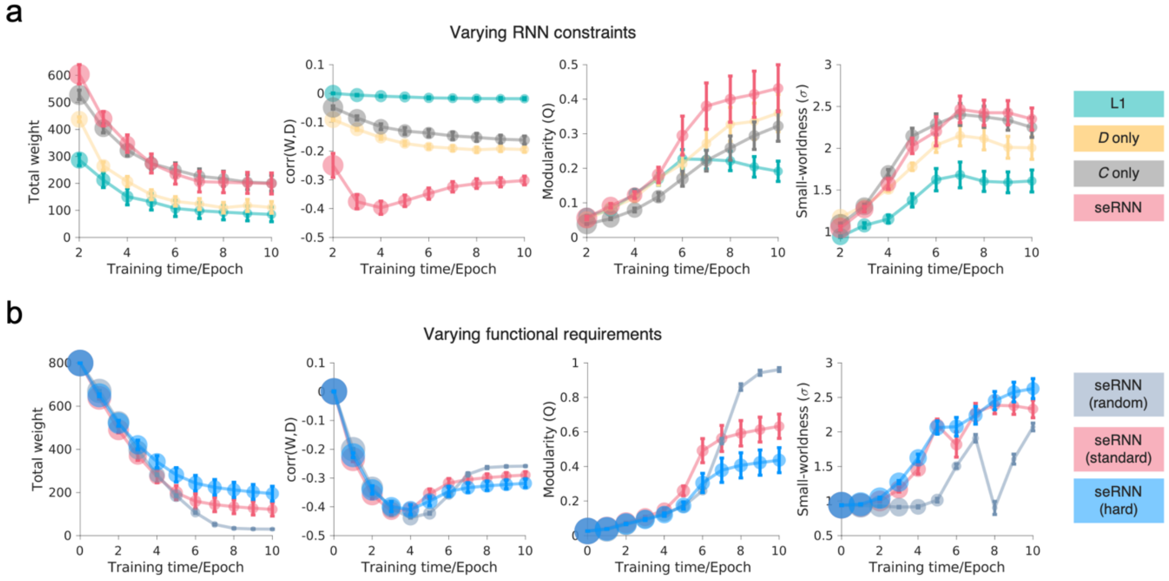


***Supplementary Figure 7.*** **Independent replication of key results across a range of baseline regularizers, including various regularization and task comparisons. a** RNNs are shown with varying regularization constraints to compare their effects on topology. L1 networks are shown in green. *D* only networks were regularized by space but not communicability and are shown in yellow. *C* only networks were regularized by communicability but not space and are shown in grey. seRNNs are the same networks that have been regularized by both and are shown in pink. These networks are all functioning and have been thresholded $> 90\%$ validation accuracy. Lines correspond to 2SEs across the 100 networks. The size of the node corresponds to the mean weights. All networks are independent to any that have been shown in the main text, and each network class was trained incrementally over 100 regularization strengths. **b** The statistical structure of the task was altered to observe the effect of various functional requirements on the topology of the resultant seRNNs. Standard seRNNs are shown in pink. seRNNs trained on a randomized task setup are shown in grey. seRNNs trained on the harder task setup are shown in blue. All networks are functioning and have been thresholded $> 90\%$ validation accuracy, apart from seRNN (random) which were non-functional networks due to the nature of the task. As above, lines correspond to 2SEs across the 100 networks. The size of the node corresponds to the mean weights. All networks are independent of any that have been shown in the main text, and each network class was trained incrementally over 100 regularization strengths.

#### ***Generative network modelling of RNN architectures***

Since Betzel, et al. (2016), the most common way of fitting simulated networks via the generative network model has been to examine its *Energy*: which is the maximum Kolmogorov-Smirnov statistics between four commonly used graph theory measures. As expanded on in the main report, homophilic generative rules – which simulate networks based upon a measure of topological self-similarity – tend to simulate networks uniquely well (Betzel, et al. 2016; Akarca, et al; 2021; Akarca, et al; 2022; Carozza, et al; 2022).

We show in **Supplementary Figure 8**, that unlike seRNNs, L1 networks cannot be uniquely simulated by homophily rules.

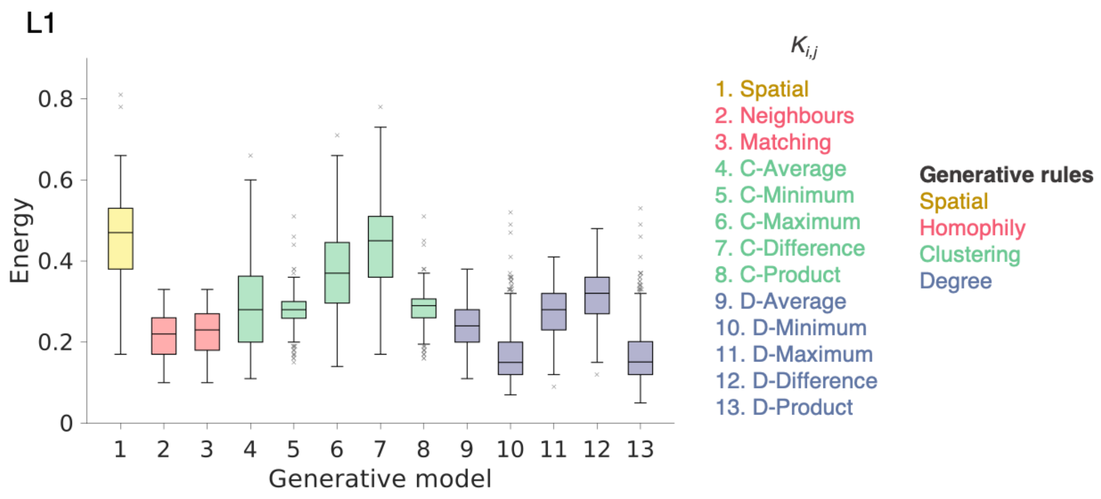


***Supplementary Figure 8*. L1 network generative model energy profile.** For L1 networks there is no single set of generative rules that generates the lowest energy. For each generative model, the best performing simulation (indexed by the energy) is shown – fit to each of the 455 functioning L1 networks.

Generative modelling findings are likely to change over time or development (Betzel, et al; 2016; Akarca, et al; 2022) due to the underlying developmental changes that occur in networks. The current framework allows us to probe the precise effect of training time and regularization (i.e., constraints) on generative model performance. If homophilic mechanisms result from spatial/communication constraints within empirical networks, we would expect two phenomena: that homophily model fits improve with both (1) training time (as more time would have elapsed for more constraints to have occurred) and (2) regularization constraints (as more constraints have been imposed).

Indeed, we find both to be the case (**Supplementary Figure 9**). First, model fits attained by homophilic models improve, relative to all other models, with more training, from epoch 3 (where homophily does worse than the mean of all other models) through to epoch 9 (where homophily performs >2x better than the mean of all other models; **Supplementary Figure 9a**). Second, we find that by correlating the energy achieved by different generative models with the regularization constraints imposed, we find that there is a unique negative correlation for homophilic rules (**Supplementary Figure 9b**).

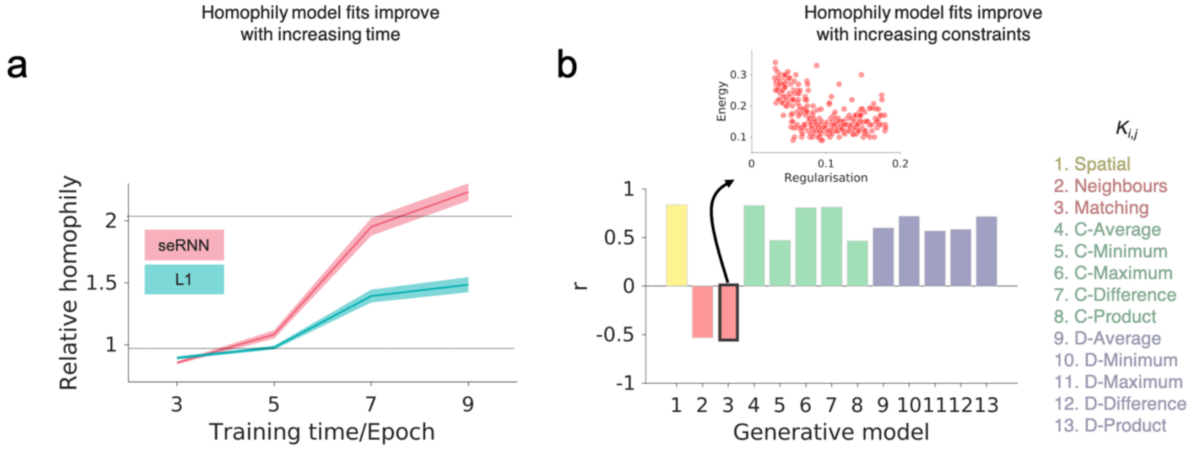


***Supplementary Figure 9.*** **Homophily model fits improve with time and constraints.**
**a** Homophily models do increasingly well over epochs/training time over the duration of training relative to all other models. The relative homophily measure is calculated as the ratio between the energy of homophily generative models relative to all other generative models. The greater the relative homophily means the better that homophily generative models are at simulating the topology of the network. **b** Networks which are regularized more are better approximated by homophily models, uniquely. Inset scatter plot shows the relationship between regularization and matching energy.

#### ***Spatial configuration of task-relevant variables***

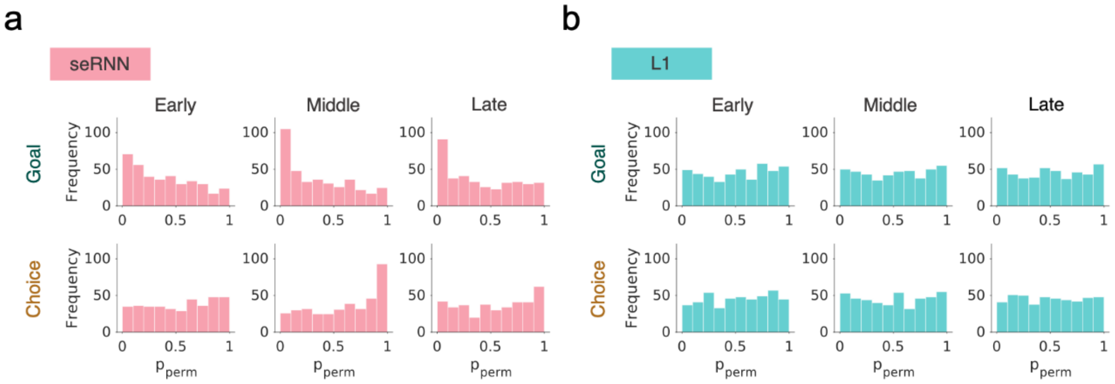


***Supplementary Figure 10.*** **Replication of spatial configurations using a weighted variation of the spatial permutation statistical procedure.** Terminology and conventions as in **Figure 3** and this procedure is outlined in **Methods;** ***Spatial permutation of task-relevant information***.

#### ***Structure-function relationships in seRNNs***

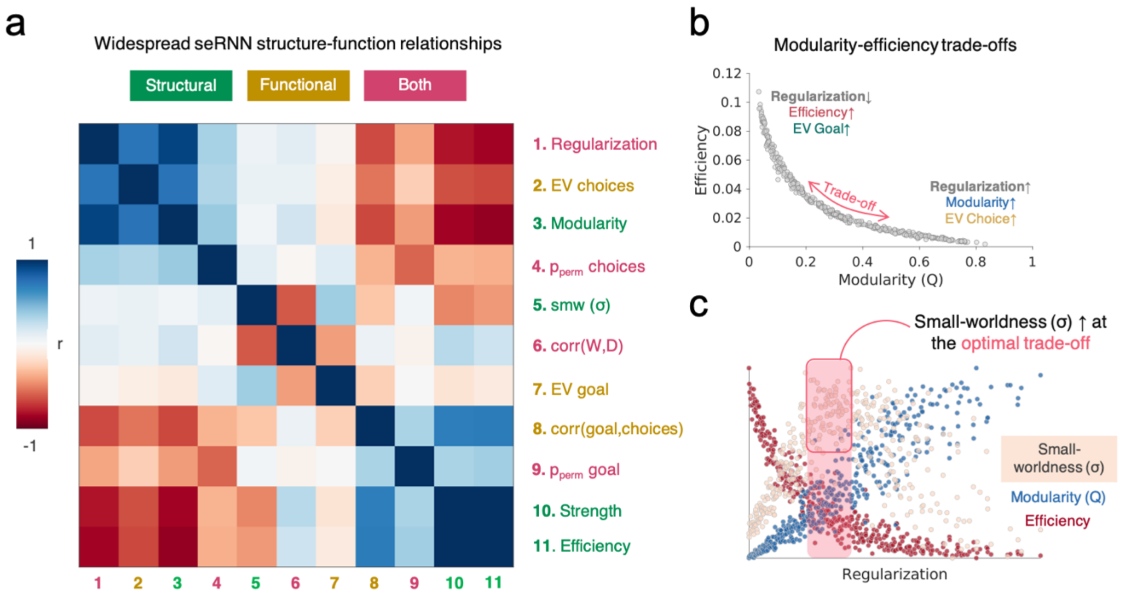


***Supplementary Figure 11.*** **Intertwined global structural-functional relationships in seRNNs. a** The correlation matrix of structural (green), functional (yellow) and both (pink) measures in the seRNNs shows wide-spread relationships that cluster. **b** A scatter plot showing a negative relationship between global efficiency versus modularity across seRNNs. Each plot in the scatter is a seRNN of a different regularization strength. seRNNs that have not been regularized as much tend to have higher global efficiency and less modularity, and sit to the left of the curve. These networks tend to have a greater explained variance for the goal. In contrast, seRNNs that have been regularized more tend to have lower global efficiency and higher modularity, and therefore sit to the right of the curve. These networks tend to have a greater explained variance for the choice. **c** A scatter plot of regularization (x-axis) versus efficiency (red; decreases), modularity (increases; blue) and small-worldness which peaks at the point at which there is a trade-off between modularity and efficiency and then declines thereafter. This inverted-U function peaks within a window of regularization that is highlighted in purple and continued into the next panel. The y-axis is normalized and therefore the scale is removed.
